# Coupled Control of Distal Axon Integrity and Somal Responses to Axonal Damage by the Palmitoyl Acyltransferase ZDHHC17

**DOI:** 10.1101/2020.09.01.276287

**Authors:** Jingwen Niu, Shaun S. Sanders, Hey-Kyeong Jeong, Sabrina M. Holland, Yue Sun, Kaitlin M. Collura, Luiselys Hernandez, Haoliang Huang, Michael R. Hayden, George M. Smith, Yang Hu, Yishi Jin, Gareth M. Thomas

**Affiliations:** Shriners Hospitals Pediatric Research Center, Lewis Katz School of Medicine at Temple University, 3500 N. Broad Street, Philadelphia, PA 19140; Section of Neurobiology, Division of Biological Sciences, University of California San Diego, La Jolla, CA 92093; Department of Ophthalmology, Stanford University School of Medicine, Palo Alto, CA, 94304; Department of Medical Genetics, Centre for Molecular Medicine and Therapeutics, The University of British Columbia, Vancouver, British Columbia, Canada; Department of Neuroscience, Lewis Katz School of Medicine at Temple University; Department of Anatomy and Cell Biology, Lewis Katz School of Medicine at Temple University

**Author notes:** Equal contribution (Jingwen Niu, Shaun S. Sanders).

## Abstract

After optic nerve crush (ONC), the cell bodies and distal axons of most retinal ganglion cells (RGCs) degenerate. RGC somal and distal axon degeneration were previously thought to be controlled by two distinct pathways, involving activation of the kinase DLK and loss of the axon survival factor NMNAT2, respectively. However, we found that mutual palmitoylation by the palmitoyl acyltransferase ZDHHC17 couples the DLK and NMNAT2 signals, which together form a “trust, but verify system”. In healthy optic nerves, ZDHHC17-dependent palmitoylation ensures NMNAT-dependent distal axon integrity, while following ONC, ZDHHC17-dependent palmitoylation is critical for DLK-dependent somal degeneration. We found that ZDHHC17 also controls survival-versus-degeneration decisions in sensory neurons and identified motifs in NMNAT2 and DLK that govern their ZDHHC17-dependent regulation. These findings suggest that the control of somal and distal axon integrity should be considered as a single, holistic process, involving two palmitoylation-dependent pathways acting in concert.

## Introduction

Responses of Retinal Ganglion Cells (RGCs) to Optic Nerve Crush (ONC) are widely studied, in part to better understand how RGCs degenerate in conditions such as glaucoma (Aguayo et al. 1991; Danesh-Meyer 2011; Hu et al. 2012; Kalesnykas et al. 2012). Understanding the coordination and control of ONC responses may also provide insights into additional forms of nervous system injury and/or other neuropathological conditions (Benowitz et al. 2017; Tran et al. 2019).

After ONC, the cell bodies and distal axons of most RGCs degenerate (Berkelaar et al. 1994; Harvey 2007; Hu et al. 2012). RGCs have limited intrinsic capacity for long distance regeneration, so this response avoids a situation in which severed RGC axons remain intact in the absence of input, or in which RGC cell bodies remain alive but disconnected to their target(s). However, it is unclear whether and how axonal and somal responses to ONC might be coupled to avoid ‘mixed messages’ of this type, particularly because ONC physically interrupts both electrical and protein-based signaling between these two subcellular locations. More knowledge of this issue could increase understanding of conditions linked to optic nerve damage and also potentially provide insights into how responses to axonal insult or injury are controlled in other neuronal types.

Most reports suggest that RGC somal and distal axon degeneration involve separate proteins and pathways. In particular, ONC-induced somal degeneration critically requires a Mitogen-activated Protein Kinase (MAPK) pathway involving Dual Leucine-zipper Kinase (DLK, a ‘MAP3K’) and DLK’s downstream target c-Jun N-terminal Kinase (JNK) (Fernandes et al. 2012; Fernandes et al. 2014; Watkins et al. 2013; Welsbie et al. 2017; Welsbie et al. 2013). The DLK-JNK pathway conveys an ONC-induced retrograde axon-to-soma signal to activate a pro-degenerative transcription program (Watkins et al. 2013). However, although knockout (KO) of DLK in mice strongly protects RGC somas from ONC-induced degeneration, DLK KO protects severed distal RGC axons minimally, if at all (Fernandes et al. 2014; Yang et al. 2015).

Conversely, a key event in ONC-induced distal axon degeneration is loss of the labile biosynthetic enzyme Nicotinamide Mononucleotide Adenylyl Tranferase-2 (NMNAT2) (Conforti et al. 2014; Yamagishi 2016). Studies from diverse neuron types suggest that continuous supply of NMNAT2 to distal axons *via* anterograde transport maintains axon integrity (Coleman and Freeman 2010; Gilley and Coleman 2010). Axonal injury interrupts this transport, leading to rapid degradation of remaining NMNAT2 in the severed distal axon and subsequent Wallerian degeneration (reviewed by (Gerdts et al. 2016)). Consistent with this model, ONC-induced Wallerian degeneration is prevented by the *Wld^S^* (Wallerian degeneration slow) spontaneous mutation, which codes for an unnatural, stable form of NMNAT2’s close paralog NMNAT1 (Fernandes et al. 2014; Lorber et al. 2012), or by a cytosolic form of NMNAT1 itself (cytoNMNAT1; (Yang et al. 2015)). ONC-induced Wallerian degeneration is also prevented by KO of SARM1 (Fernandes et al. 2018; Yang et al. 2015), an axonal ‘executioner’ enzyme whose activation likely lies downstream of NMNAT2 loss (Gilley et al. 2015; Gilley et al. 2017). However, neither *Wld^S^* nor SARM1 KO prevents ONC-induced RGC somal degeneration (Beirowski et al. 2008; Fernandes et al. 2014; Fernandes et al. 2018; Lorber et al. 2012).

The picture that emerges from these studies is that the DLK-JNK and NMNAT2-SARM1 pathways are largely independent in RGCs. However, we reasoned that these two pathways might be more coordinated than is initially apparent. We based this hypothesis on findings that both DLK and NMNAT2 localize to axonal transport vesicles, and that their targeting to vesicles involves the same mechanism, namely the protein-lipid modification palmitoylation (Holland et al. 2016; Milde et al. 2013). Palmitoylation is well known to facilitate trafficking of proteins in axons (Holland and Thomas 2017; Kanaani et al. 2004; Milde et al. 2013; Tortosa et al. 2017). However, the importance of palmitoylation of DLK and NMNAT2 in the optic nerve *in vivo* has not been addressed, nor has the Palmitoyl Acyltransferase (PAT)(s) responsible been identified. Addressing these questions could provide new information about these important pro-degenerative and pro-survival pathways, while also yielding insights into mechanisms that couple responses of neuronal cell bodies and distal axons to injury and insult.

Here, we use an AAV-mediated molecular replacement approach to define the importance of DLK and NMNAT2 palmitoylation for RGC somal and distal axon integrity *in vivo*. We first report that palmitoylation of DLK is critical for ONC-induced retrograde injury signaling and subsequent RGC somal degeneration. We then identify the PAT ZDHHC17 as an evolutionarily conserved regulator of DLK palmitoylation and axon-to-soma pro-degenerative signaling. We further show that DLK’s palmitoyl-site is highly homologous to that of NMNAT2 and provide evidence that ZDHHC17-dependent palmitoylation of NMNAT2 is essential for distal axon integrity, both in cultured sensory Dorsal Root Ganglion (DRG) neurons and in RGCs *in vivo*.

These findings suggest that, rather than controlling somal and axon degeneration separately, RGCs instead use ZDHHC17 to establish a coupled ‘trust, but verify’ system; ZDHHC17 supplies palmitoyl-NMNAT2 to distal axons to signal that all is well, but simultaneously supplies palmitoyl-DLK to act as a surveillance factor for axonal injury or insult. After ONC, palmitoyl-DLK-dependent retrograde signals cause somal degeneration, while interruption of palmitoyl-NMNAT2 anterograde transport triggers distal axon degeneration. This palmitoylation-dependent mechanism allows neurons to monitor and respond to changes in axonal health and coordinate responses in both distal axons and neuronal cell bodies.

## Results

### Palmitoyl-DLK is essential for Somal Responses to Axonal Injury *in vivo*

Our overall goal was to determine whether mutual palmitoylation of DLK and NMNAT2 might facilitate coupled control of axonal and somal integrity in RGCs, and potentially also in other neurons. To address these questions, we took advantage of the tractability of RGCs to genetic manipulation following intravitreal delivery of Adeno-Associated Virus (AAV; (Martin et al. 2002)). Intravitreal injection of AAV expressing Cre recombinase or shRNA allows KO or knock down of a gene of interest in RGCs, respectively, facilitating determination of the importance of that gene’s protein product. Co-injection of an shRNA-expressing AAV with a second AAV expressing an shRNA-resistant form of the wild type (wt) protein, or its corresponding palmitoyl-site mutant, allows determination of the importance of a specific palmitoylation event *in vivo* (Fig 1A).

**Figure 1:**
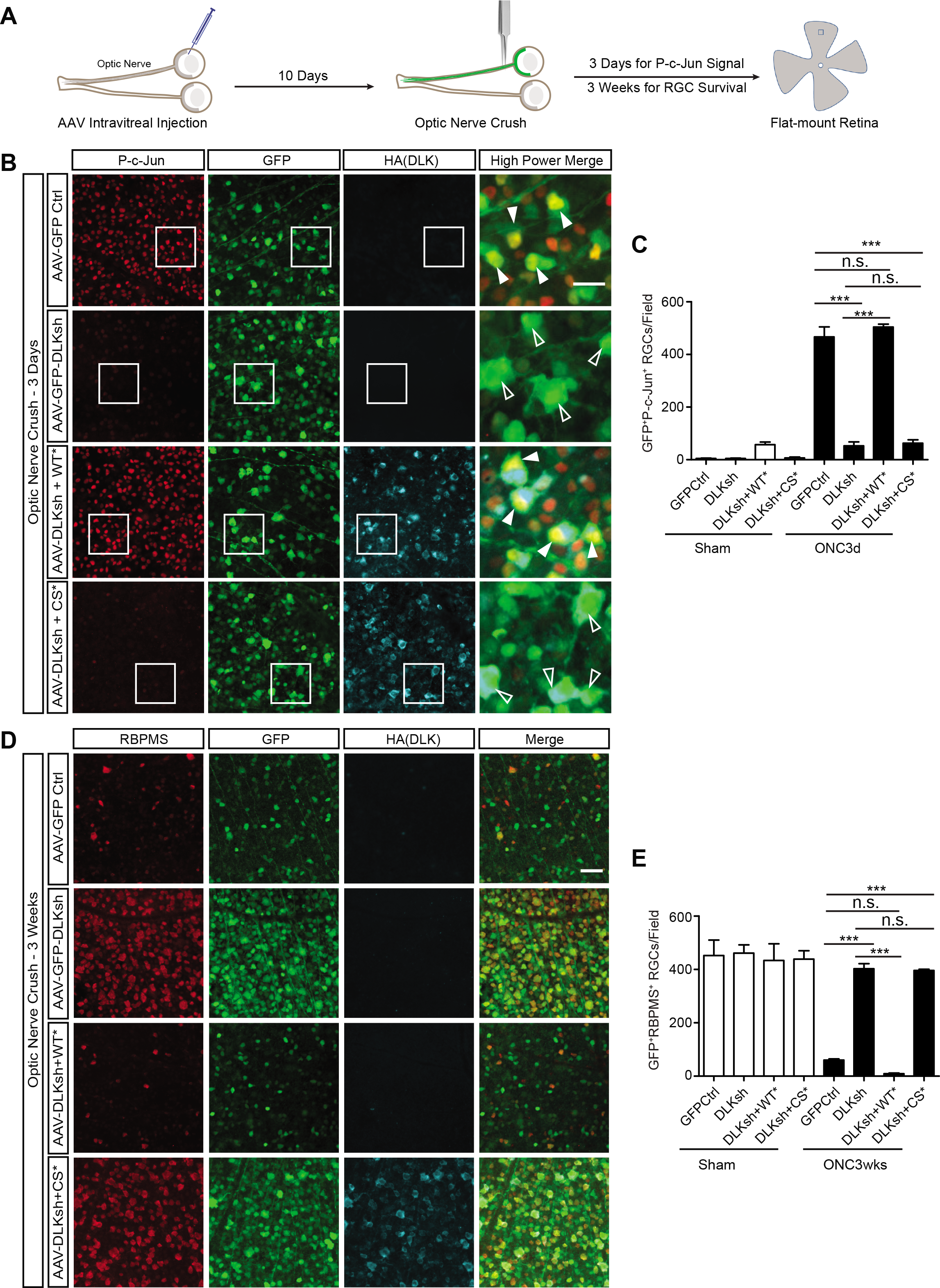
Palmitoyl-DLK is Essential for Retrograde Signaling After Optic Nerve Crush. ***A:*** Experimental set-up: adult mice were intravitreally injected with AAVs and Optic Nerve Crush (ONC) was performed 10 days later. Retinas were fixed and flat-mounted prior to immunostaining. Blue square indicates approx. region of retina from which images were acquired. ***B***: Retinal flat-mount images from eyes injected with the indicated AAVs and immunostained 3 days post-ONC with the indicated antibodies. ONC-induced P-c-Jun signal is markedly attenuated in retinas injected with DLKsh-expressing AAV (AAV-GFP-DLKsh) and ‘rescued’ by shRNA resistant wild type DLK (AAV-DLKsh + wtDLK*-HA (‘WT*’)) but not by shRNA resistant palmitoyl-mutant DLK (AAV-DLKsh + DLKCS*-HA (‘CS*’)). *4th column*: magnified merged images of boxed areas in columns 1-3. *Filled arrowheads*: phospho-c-Jun-positive AAV-infected cells. *Empty arrowheads*: phospho-c-Jun-negative AAV-infected cells. Scale bar: 20 μm ***C:*** Quantified data from *B* (n=3-5 per condition); ***: p<0.001; n.s., nonsignificant. One-way ANOVA with Bonferroni *post hoc* test. ***D:*** Retinal flat-mount images from eyes injected with the indicated AAVs, fixed 3 weeks post-ONC and immunostained with the indicated antibodies. Little HA signal remains in the AAV-DLKsh + wtDLK*-HA (‘WT*’) condition because these cells have degenerated, but extensive HA signal is still present in the AAV-DLKsh+DLKCS*-HA (‘CS*’) condition. *4th column*: Merged images from columns 1-3. Scale bar: 50 μm. ***E:*** Quantified data from *D* (n=3-5 per condition). ***: p<0.001, n.s., non-significant. Oneway ANOVA with Bonferroni *post hoc* test. All data are mean ± SEM.

We first sought to use this AAV-mediated molecular replacement approach to determine the importance of DLK palmitoylation for somal responses to ONC. In wt mice uninjected with AAV, ONC greatly increased phosphorylation of the transcription factor c-Jun in the majority of RGCs, and concomitantly decreased levels of Brn3a, a marker of healthy RGCs, consistent with prior studies (Fig S1A; (Fernandes et al. 2014; Watkins et al. 2013; Welsbie et al. 2017; Welsbie et al. 2013)). ONC also robustly increased c-Jun phosphorylation in retinas that had been intravitreally infected with control AAV (expressing GFP alone)(Fig 1B, C). ONC-induced c-Jun phosphorylation was greatly attenuated in retinas infected with AAV expressing GFP plus DLK shRNA (AAV-GFP-DLKsh; Fig 1B, C), similar to prior reports (Watkins et al. 2013; Welsbie et al. 2013). AAV-GFP-DLKsh also prevented ONC-induced increases in endogenous retinal DLK levels (Fig S1B), consistent with prior work (Watkins et al. 2013; Welsbie et al. 2013).

To define the importance of DLK palmitoylation in somal responses to ONC, we intravitreally co-delivered a second AAV to create the *in vivo* molecular replacement system described above. Co-infection of AAV-GFP-DLKsh with an AAV expressing an shRNA-resistant HA-tagged form of wtDLK (AAV-wtDLK*-HA; ‘WT*’ on Fig. 1B) restored ONC-induced c-Jun phosphorylation to wt levels (Fig. 1B, C). In marked contrast, co-infection of AAV-GFP-DLKsh plus AAV expressing shRNA-resistant HA-tagged palmitoyl-mutant (Cys127Ser mutant) DLK (AAV-DLK-CS*-HA; ‘CS*’ on Fig.1B, C) failed to rescue ONC-induced c-Jun phosphorylation (Fig. 1B, C). WtDLK*-HA and DLK-CS*-HA expressed at similar levels (Fig. 1B) and each was co-expressed in a similar fraction of GFP-expressing neurons (>80%, Fig S1C). These findings suggest that RGC somal responses to axonal injury *in vivo* require palmitoyl-DLK.

To address whether DLK palmitoylation is also critical for ONC-induced RGC somal degeneration, we used a similar dual-AAV approach but analyzed RGC viability 3 weeks post-ONC (Fig. 1A, D). Retinas were then fixed and stained to detect RBPMS, a marker whose expression in RGCs is unaffected by injury (Kwong et al. 2011). In AAV-GFP-infected retinas, ONC dramatically reduced RGC numbers (Fig 1D, E), consistent with prior studies (Fernandes et al. 2014; Watkins et al. 2013; Welsbie et al. 2013). However, in retinas infected with AAV-GFP-DLKsh, ONC did not reduce RGC numbers (Fig 1D, E). ONC-induced RGC loss was fully rescued by AAV-wtDLK*-HA, but not by AAV-DLK-CS*-HA (Fig 1D, E; ‘WT*’, ‘CS*’ on Fig. 1D, E). These findings suggest that palmitoyl-DLK is essential for ONC-induced RGC somal degeneration.

### The Palmitoyl Acyltransferase (PAT) ZDHHC17 is an evolutionarily conserved regulator of DLK

Having defined the functional importance of DLK palmitoylation in somal responses to ONC, we next sought to identify the PAT(s) responsible. RGCs express >20 of the 24 mammalian PATs (Sajgo et al. 2017; Tran et al. 2019), potentially complicating this task. However, we realized that this search could be narrowed by our finding that localization of the *C. elegans* DLK ortholog, CeDLK-1, is highly palmitoylation-dependent in worm sensory neurons (Holland et al. 2016). This result suggested that DLK palmitoylation is highly evolutionarily conserved and that the DLK PAT might thus also be a conserved enzyme. Of several PATs that palmitoylate cotransfected mammalian DLK (Holland et al. 2016), the most evolutionarily conserved is ZDHHC17 (Roth et al. 2002; Young et al. 2012). In contrast to most PATs, ZDHHC17 possesses six, rather then four, predicted Transmembrane Domains (TMDs) in addition to the enzymatic DHHC domain, plus an N-terminal Ankyrin repeat (AnkR) region (Fig 2A). Two *C. elegans* presumptive ZDHHC17 orthologs, *dhhc-13* and *dhhc-14*, possess a similar TMD arrangement and AnkR region (Edmonds and Morgan 2014)(Fig 2A). When we examined GFP-tagged DLK-1 localization in *C. elegans* mechanosensory axons, we found that in worms carrying null mutants for *dhhc-13* and *dhhc-14*, GFP-CeDLK-1 shifted from a punctate to a more diffuse localization (Fig 2B, C, mutants described in Fig S2A). These findings suggest that ZDHHC17 orthologs regulate CeDLK-1.

**Figure 2:**
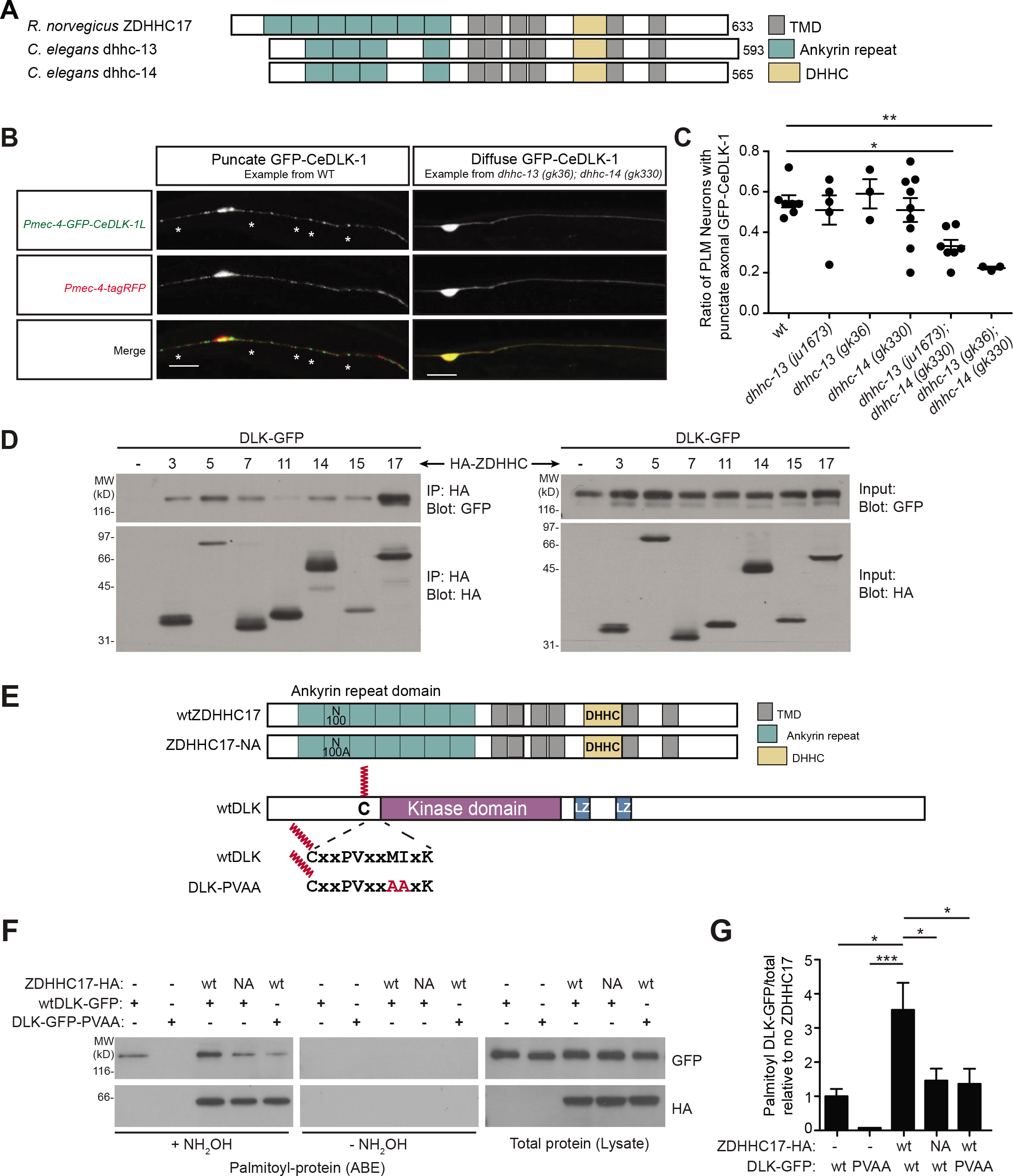
ZDHHC17 is an Evolutionarily Conserved Regulator of DLK Palmitoylation. ***A:*** Schematic of Rat ZDHHC17 aligned with Ce-dhhc13 and Ce-dhhc14, the only *C. elegans* PATs containing N-terminal ankyrin repeats (Edmonds and Morgan 2014). The number of transmembrane domains (TMDs) for Ce-dhhc-13 and Ce-dhhc-14 is also similar to ZDHHC17 but not other mammalian PATs. Catalytic DHHC domain is also shown. ***B:*** Images of sensory neurons expressing GFP-tagged CeDLK-1 (GFP-CeDLK-1) (*top*) and cell fill tagRFP (*middle*) from wt and *dhhc-13/dhhc-14* double mutant worms. *Bottom*: merged images of GFP-CeDLK-1 and TagRFP signals. GFP-CeDLK-1L is punctate along axons and in the soma of most wt PLM neurons, consistent with (Holland et al. 2016). GFP-CeDLK-1L puncta are less frequently observed in PLM of *dhhc-13(ju1673); dhhc-14(gk330*) and *dhhc-13(gk36); dhhc-14(gk330*) mutants. Scale bar: 20 μm. ***C:** Ce dhhc-13/dhhc-14* mutation significantly reduces punctate GFP-CeDLK-1 localization. Each data point represents the ratio of PLM with punctate GFP-CeDLK-1 to total, from an independent group of 30-70 animals. **: p<0.01, *:p<0.05 relative to wt, one way ANOVA. ***D:*** HA immunoprecipitates (IPs, *left*) and parent lysates (*right*) from HEK293T cells transfected to express GFP-tagged wtDLK (DLK-GFP) plus the indicated HA-tagged ZDHHC-PATs, blotted with the indicated antibodies. Similar results were obtained in two other experiments. ***E:*** ZDHHC17 palmitoylation of DLK involves an AnkR-zDABM interaction. *Upper schematic*: ZDHHC17 domain arrangement, showing N100A (ZDHHC17-NA) mutation that disrupts binding to zDABM-containing substrates (Verardi et al. 2017). *Lower schematic*: DLK domain arrangement showing PV--AA mutation that disrupts zDABM-like motif. ***F:*** Acyl-Biotinyl Exchange (ABE) samples (*left*) and parent lysates (*right*) from HEK293T cells transfected to express the indicated forms of DLK-GFP and HA-ZDHHC17, blotted with the indicated antibodies. *Center*: signals from parallel ABE assays lacking the key reagent hydroxylamine (NH2OH). ZDHHC17-N100A mutation and DLK-GFP PV--AA mutation both reduce DLK-GFP palmitoylation by ZDHHC17. ***G:*** Quantified DLK-GFP palmitoylation from multiple determinations from *F*. (N=4); *:p<0.05, ***:p<0.001 relative to wild type, one-way ANOVA with Bonferroni *post hoc* test.

### Preferential Regulation of DLK by ZDHHC17 *via* AnkR-mediated Binding

We next addressed whether ZDHHC17 is a PAT for mammalian DLK. In cotransfected HEK293T cells, ZDHHC17 coimmunoprecipitated GFP-tagged wtDLK (wtDLK-GFP) more effectively than did six other PATs that can robustly palmitoylate cotransfected DLK (Holland et al. 2016)(Fig 2D). Of theses PATs, only ZDHHC17 contains an AnkR, suggesting ZDHHC17 might use this region to bind DLK. ZDHHC17’s AnkR was reported to recognize a ZDHHC Ankyrin Domain Binding Motif (zDABM) found in several ZDHHC17 substrates (Lemonidis et al. 2015). DLK lacks the strict consensus zDABM sequence [IV][IV]XXQP (where X is any amino acid)(Lemonidis et al. 2015), but contains a sequence PVXXMIXK adjacent to its palmitoylation site (Fig 2E). This sequence matches a subsequently reported broader zDABM consensus [AIPV][ITV]XX[CMQ][IP]X[KR] (Lemonidis et al. 2017).To test whether DLK binds ZDHHC17 *via* this region, we expressed a point mutant of this DLK zDABM (PV--AA mutant; Fig 2E, S2B) and observed markedly reduced localization of DLK-GFP to Golgi membranes, which is known to be palmitoylation-dependent (Fig S2C) (Martin et al. 2019). Golgi enrichment of DLK-GFP was increased by cotransfected wt HA-tagged ZDHHC17 (wt-ZDHHC17-HA), but was reduced in cells co-expressing HA-tagged ZDHHC17-N100A, an AnkR mutant that fails to recognize zDABM-containing substrates ((Verardi et al. 2017) ‘ZDHHC17-HA NA’, Fig S2B, C, schematic on Fig 2E). DLK PV--AA and ZDHHC17-NA mutations also both reduced palmitoylation of DLK-GFP by cotransfected ZDHHC17 in biochemical assays (Fig 2F, G). Together, these findings suggest that DLK is preferentially recognized by ZDHHC17 compared to other PATs and that DLK palmitoylation by ZDHHC17 involves an AnkR-zDABM interaction.

### ZDHHC17 controls DLK palmitoylation and signaling in mammalian neurons in culture and *in vivo*

Like DLK, ZDHHC17 is neuronally-enriched (Fagerberg et al. 2014; Yue et al. 2014) and is also one of the most highly expressed PATs in RGCs (Sajgo et al. 2017; Tran et al. 2019). These findings suggest that ZDHHC17 is thus a strong candidate to palmitoylate endogenous DLK. However, the retina is not well suited to biochemical palmitoylation assays, so we addressed this question using cultured DRG neurons, in which somal responses to axonal insult and injury also require DLK and its palmitoylation (Ghosh et al. 2011; Holland et al. 2016; Simon et al. 2016). Lentiviral infection of DRG neurons with either of two *Zdhhc17* shRNAs greatly reduced DLK palmitoylation but only minimally affected DLK protein levels (Fig 3A, B, Fig S3A) and did not affect palmitoylation of another axonal palmitoyl-protein, GAP43 (Fig 3A, B). In contrast to the marked effect of ZDHHC17 knockdown, palmitoyl-DLK levels were unaffected by combined knockdown of ZDHHC5 and ZDHHC8, two PATs that strongly palmitoylate DLK in non-neuronal cells and that often compensate for one another in neurons (He et al. 2014; Holland et al. 2016; Thomas et al. 2012) (Fig S3B). These results suggest that ZDHHC17 is a major regulator of DLK palmitoylation in DRG neurons.

**Figure 3:**
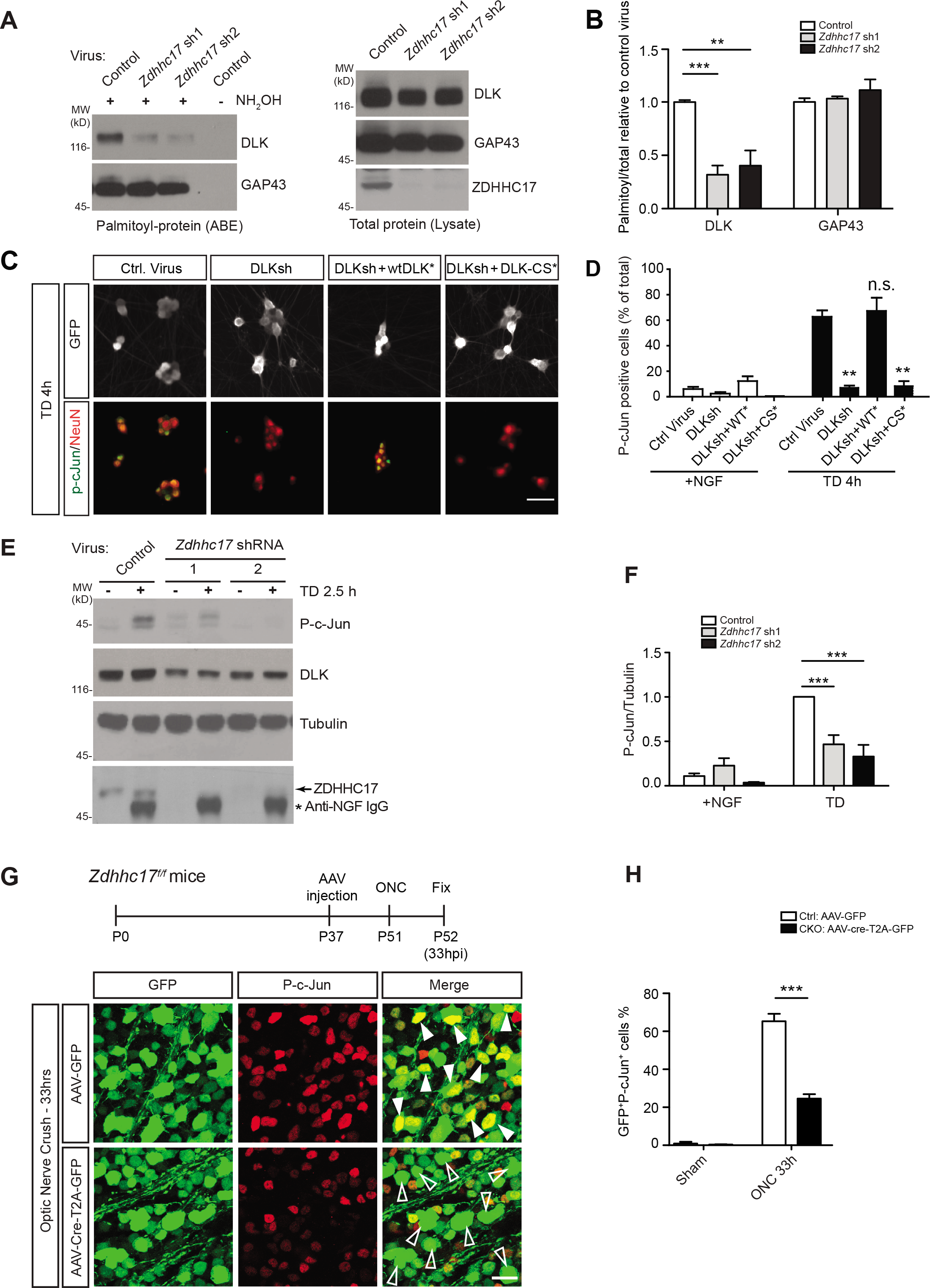
ZDHHC17 Controls Neuronal DLK Palmitoylation and Signaling *in vitro* and *in vivo*. ***A:*** DRG neurons were infected with the indicated lentiviruses and palmitoyl-proteins were purified by ABE. Palmitoyl-DLK and -GAP43 are present in ABE fractions from cultures infected with control virus and absent from a control lacking hydroxylamine (NH2OH). *Zdhhc17* knockdown by either of two shRNAs (confirmed on *bottom right panel* and Fig. S3A) greatly reduces DLK, but not GAP43, palmitoylation. ***B***: Histograms of quantified palmitoyl:total ratios for DLK and GAP43, n=5 individual cultures; ***:p<0.001, **:p<0.01 versus control virus condition, One-way ANOVA with Bonferroni *post hoc* test. ***C:*** Images of GFP signal (top) and merged p-cJun and NeuN signals (bottom) from cultured DRG neurons infected with the indicated lentiviruses, subjected to TD for 4h and immunostained. Scale bar: 50 μm. DLK knockdown greatly reduces TD-induced c-Jun phosphorylation. ShRNA-resistant wtDLK (wtDLK*), but not shRNA-resistant DLK-CS (DLK-CS*), rescues this effect. ***D:*** Quantified data from *C*. **P<0.01 versus control virus TD condition. One-way ANOVA with Bonferroni *post hoc* test, n= 3-6 determinations per condition. ***E:*** Lysates from cultured sensory neurons infected with control virus (Control) or virus expressing either of two different *Zdhhc17* shRNAs, subjected to TD for 2.5 h or left unstimulated, lysed and blotted with the indicated antibodies. Lower panel shows ZDHHC17 (arrowed) and anti-NGF antibody used during TD, detected by secondary antibody (asterisk). ***F:*** Quantified data from *E* confirms reduced TD-induced c-Jun phosphorylation in *Zdhhc17* knockdown neurons. n=5; ***:p<0.001 versus control virus condition, 2-way ANOVA with Bonferroni *post hoc* test (virus p=0.0003, TD p<0.0001, interaction p=0.0003). ***G:** Zdhhc17* conditional knockout in retina blocks DLK-dependent signaling. *Top*: Experimental timeline: *Bottom*: Images of flat-mounted retinas from *Zddhc17^f/f^* mice injected with the indicated AAVs, subjected to ONC, fixed 33h later and immunostained with the indicated antibodies. *Right panels*: merged images from columns 1-2. *Filled arrowheads*: phospho-c-Jun-positive AAV-infected cells. *Empty arrowheads*: phospho-c-Jun-negative AAV-infected cells. Scale bar: 30 μm. ***H:*** Histogram of numbers of AAV-GFP- and AAV-Cre-T2A-GFP-infected neurons that are also positive for p-cJun. *Zdhhc17* conditional knockout (CKO) reduces ONC-induced c-Jun phosphorylation. ***; p<0.001, one-way ANOVA with Bonferroni *post hoc* test, n=4 determinations per condition. All data are mean ± SEM.

### DLK-dependent Somal Responses to Axonal Insult and Injury Require ZDHHC17

We next asked whether loss of ZDHHC17 phenocopies loss of palmitoyl-DLK to prevent somal responses to axonal insult or injury. In cultured DRG neurons, the somal response to Trophic factor Deprivation (TD) is strongly DLK-dependent (Ghosh et al. 2011; Simon et al. 2016). Multiple insights into molecular regulation of DLK signaling have been gained by combining this TD model with ONC studies (Huntwork-Rodriguez et al. 2013; Larhammar et al. 2017; Simon et al. 2016; Yang et al. 2015). We therefore asked whether the DRG somal response to TD also requires palmitoyl-DLK. Acute removal of the neurotrophin Nerve Growth Factor (NGF) triggered robust c-Jun phosphorylation in DRG neuron somas, consistent with prior studies (Ghosh et al. 2011; Simon et al. 2016). As in RGCs subjected to ONC, cJun phosphorylation was prevented by DLKshRNA and rescued by shRNA-resistant wt, but not palmitoyl-mutant, DLK (wtDLK*, DLK-CS*; Fig 3C, D). This result suggested that, as in injured RGCs (Fig. 1), cJun phosphorylation in trophically-deprived DRG neurons requires palmitoyl-DLK.

To determine whether this reduced somal response to TD in the absence of palmitoyl-DLK is phenocopied by loss of ZDHHC17, we infected DRG neurons with control lentivirus, or with lentivirus expressing either of our two *Zdhhc17* shRNAs, and subsequently performed TD. Consistent with our hypothesis, *Zdhhc17* knockdown greatly reduced TD-induced c-Jun phosphorylation (Fig 3E, F).

To address whether loss of ZDHHC17 phenocopies loss of palmitoyl-DLK *in vivo*, we intravitreally infected *Zdhhc17^f/f^* mice (Sanders et al. 2016) with AAV-Cre-GFP (expressing Cre recombinase linked *via* a self-cleaving T2A sequence to GFP) or with control AAV-GFP. We performed ONC nine days post-AAV injection and subsequently fixed retinas to assess c-Jun phosphorylation (Fig 3G). Phospho-c-Jun/GFP double-positive cells were readily observed in AAV-GFP-infected retinas after ONC (Fig 3G, H) but were far less detectable in AAV-Cre-GFP-injected (i.e. *Zdhhc17* conditional KO (CKO)) retinas (Fig 3G, H). These findings support the hypothesis that ZDHHC17 palmitoylates DLK and suggest that this process is essential for somal responses to axonal injury/insult in both DRG neurons and RGCs.

We also asked whether ZDHHC17 is a likely DLK PAT in other nervous system cell types. Data extraction from an online single cell RNA-Seq resource (www.mousebrain.org, compiled using data from (Zeisel et al. 2018) and related studies) revealed that DLK (gene name *Map3k12*) mRNA levels correlate better with *Zdhhc17* mRNA levels than with mRNA levels of any other PAT across 265 nervous system cell types (Fig S3C, D). ZDHHC17 is thus highly expressed in cell types that strongly express DLK, further supporting the hypothesis that ZDHHC17 is a major PAT for DLK.

### ZDHHC17 also binds and palmitoylates NMNAT2 in heterologous cells and in neurons

Having identified ZDHHC17 as a likely PAT for DLK, we asked whether ZDHHC17 is also a major PAT for NMNAT2. Several lines of evidence supported this hypothesis. First, the regions surrounding the DLK and NMNAT2 palmitoyl-sites are highly homologous - indeed, bioinformatic search tools (Blanc et al. 2015; Obenauer et al. 2003) revealed that these regions are more similar to one another than to any other known palmitoyl-site (Fig 4A). Second, NMNAT2 also contains a bioinformatically predicted zDABM close to its palmitoyl-motif (Lemonidis et al. 2017)(Fig 4B). Both the palmitoyl-site region and the zDABM are highly conserved in all vertebrate NMNAT2 orthologs (Fig 4B), although the functional importance of NMNAT2’s zDABM has not been previously addressed in any system. In addition, ZDHHC17 was previously implicated as a potential PAT for NMNAT2, albeit based largely on findings from non-neuronal cells (Milde and Coleman 2014).

**Figure 4:**
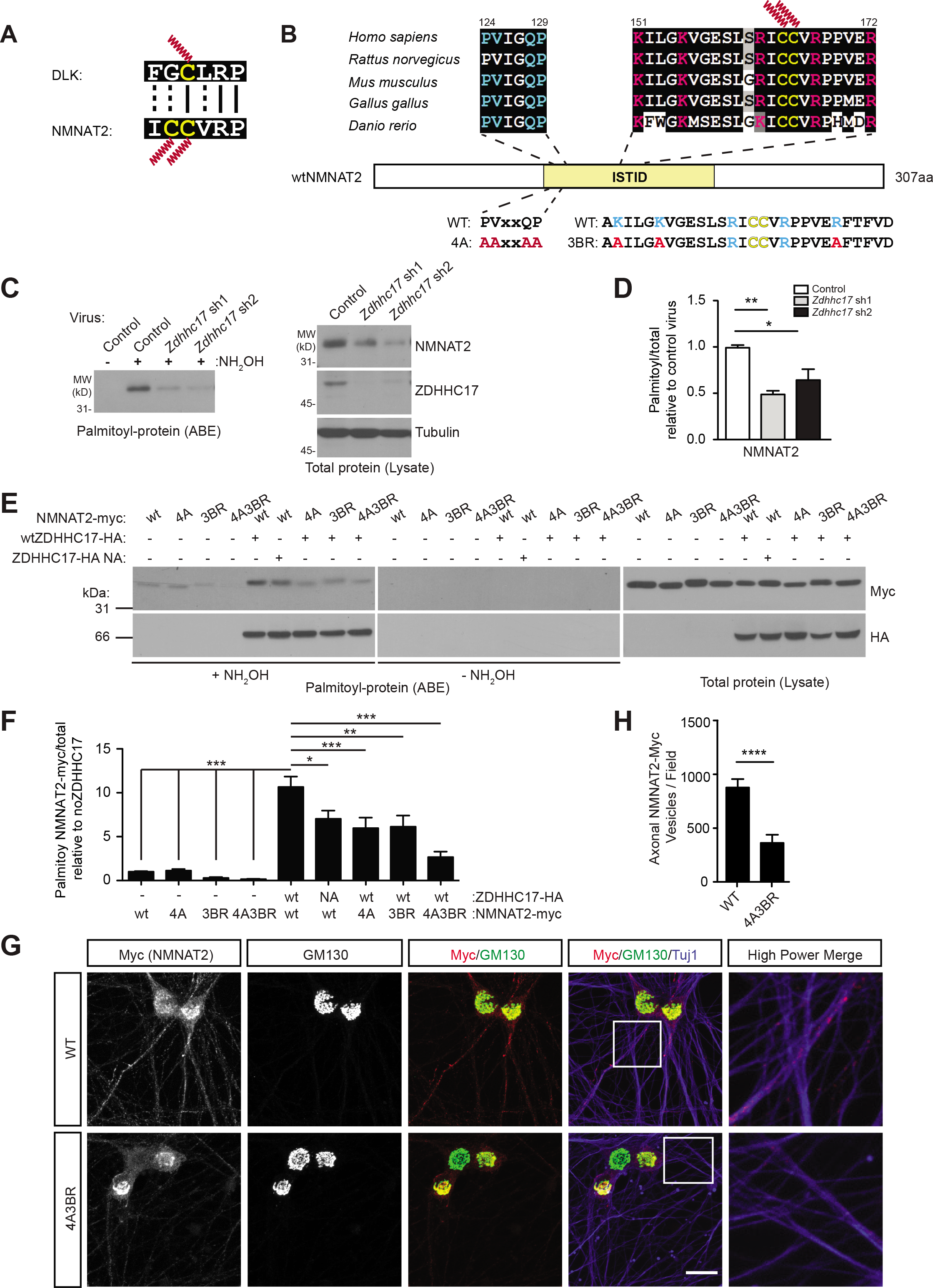
DLK and NMNAT2 Palmitoyl-sites are highly similar and NMNAT2 Palmitoylation and Localization is also ZDHHC17-dependent. ***A:*** Sequences around DLK (top) and NMNAT2 palmitoyl-sites (bottom) are highly homologous. DLK and NMNAT2 regions shown are 100% conserved in all vertebrate species examined. Palmitoyl-cysteines are shown in yellow, along with red palmitate lipid. ***B:*** NMNAT2’s central Isoform Specific Targeting and Interaction Domain (ISTID, yellow) contains multiple conserved elements linked to palmitoylation-dependent regulation: *cyan*: zDABM consensus; *magenta*: basic residues (BR) important for membrane attachment and palmitoylation (Milde et al. 2013); *yellow*: palmitoyl-cysteines. zDABM and BR residues mutated in NMNAT2-4A, -3BR and -4A-3BR combination mutants are shown below. ***C:** Zdhhc17* knockdown reduces NMNAT2 palmitoylation in neurons. ABE and total lysate samples from DRG sensory neurons infected with the indicated lentiviruses were immunoblotted to detect total and palmitoyl-NMNAT2. Palmitoyl-NMNAT2 is strongly detected in ABE fractions from cultures infected with control virus and absent from a control lacking hydroxylamine (NH2OH). *Zdhhc17* knockdown using either of two shRNAs (confirmed by western blot, *middle right*) greatly reduces endogenous NMNAT2 palmitoylation. ***D:*** Histograms of palmitoyl:total ratios for NMNAT2, from n=5 individual cultures. **:p<0.01, *:p<0.05 versus control virus condition, One-way ANOVA with Bonferroni *post hoc* test. ***E:*** NMNAT2 palmitoylation by ZDHHC17 involves an AnkR-zDABM interaction. Western blots of ABE samples (*left*) and parent lysates (*right*) from HEK293T lysates expressing the indicated forms of ZDHHC17-HA and NMNAT2-myc. NMNAT2 palmitoylation is reduced by ZDHHC17-N100A mutation, or by NMNAT2-4A and 3BR mutations individually, and is reduced further in the NMNAT2-4A-3BR combination mutant. ***F:*** Quantified data from n=7 determinations per condition from *E*. *<0.05, **<0.01, ***<0.001; one-way ANOVA with Bonferroni *post hoc* test. ***G:*** Images of DRG neurons lentivirally infected to express myc-tagged NMNAT2wt or 4A-3BR mutant, immunostained with the indicated antibodies. *Right hand column*: magnified view of the boxed area in the fourth column images. Scale bar: 15 μm. ***H:*** Quantified data from n=3 determinations per condition from *G* confirms decreased axonal targeting of NMNAT2-4A-3BR mutant. ****; p<0.001, t-test.

We verified that ZDHHC17 robustly bound and palmitoylated cotransfected NMNAT2 (Fig S4A, B), consistent with this prior report (Milde and Coleman 2014). In addition, both *Zdhhc17* shRNAs markedly reduced palmitoyl-NMNAT2 signals in DRG neuron ABE fractions and also, to a lesser extent, reduced NMNAT2 total expression (Fig 4C, D). These findings suggest that ZDHHC17 is a major NMNAT2 PAT in transfected cells and in neurons.

We next assessed the extent to which NMNAT2’s zDABM is important for palmitoylation by ZDHHC17. Interestingly, mutation of the zDABM alone (NMNAT2-4A mutant) partially reduced NMNAT2 palmitoylation by ZDHHC17 (Fig 4E, F). Mutation of ZDHHC17’s critical asparagine that is required to recognize zDABM-containing substrates (N100A mutation) also significantly, though only partially, reduced palmitoylation of NMNAT2 by ZDHHC17 (Fig 4E, F). However, another NMNAT2 region that is important for its palmitoylation is a group of 5 basic residues surrounding the palmitoyl-motif. Combined mutation of these 5 basic residues (ΔBR mutant) was previously reported to markedly reduce NMNAT2 palmitoylation (Milde et al. 2013). Importantly, though, two of these five sites (R162, R167) lie very close to the sites of NMNAT2 palmitoylation (C164, C165; Fig 4B) and it is possible that their mutation affects NMNAT2 recognition by the NMNAT2 PAT. We therefore assessed palmitoylation of a more subtle NMNAT2-3BR mutant (containing only 3 of the 5 basic site mutations, most distal from NMNAT2’s palmitoyl-sites) by ZDHHC17. Like the NMNAT2-4A mutant, the NMNAT2-3BR mutant was also less well palmitoylated than NMNAT2wt (Fig 4E, F), and combining this 3BR mutation with the zDABM mutation (NMNAT2-4A-3BR) further reduced NMNAT2 palmitoylation by ZDHHC17 (Fig 4E, F). These results suggest that DLK and NMNAT2 not only share similar palmitoyl-sites, but also both possess motifs that facilitate their palmitoiylation by ZDHHC17. While either the zDABM or BR regions in isolation are largely sufficient to allow NMNAT2 palmitoylation, mutation of both regions markedly impairs palmitoylation of NMNAT2.

To assess whether the zDABM and/or BR region are important for NMNAT2 targeting in neurons, we infected neurons with lentivirus expressing either wt or 4A-3BR variants of NMNAT2-myc. Consistent with prior reports (Berger et al. 2005; Milde et al. 2013), NMNAT2-wt-myc colocalized with the Golgi marker GM130 and was also detected on axonal puncta that are likely vesicles (Fig 4G). In contrast, NMNAT2-4A-3BR-myc still colocalized with GM130, but was much less readily detected in axons (Fig 4G, H). This lack of detection was not due to reduced expression because wtNMNAT2-myc and NMNAT2-4A3BR-myc protein levels in infected cultures were very similar (Fig S4C, D). These findings suggest that, as in non-neuronal cells (Fig 4E, F), the combination of the BR and zDABM motifs control neuronal NMNAT2 palmitoylation, which is in turn important for NMNAT2 axonal targeting.

We also asked whether ZDHHC17 likely regulates NMNAT2 in other nervous system cell types. As with DLK, extracted RNA-Seq data revealed that *Nmnat2* mRNA expression correlate better with *Zdhhc17* mRNA levels than with mRNA levels of any other PAT across 265 nervous system cell types (Fig S4E, F). Together, these findings suggest that NMNAT2 is indeed an endogenous ZDHHC17 substrate in DRG sensory neurons and perhaps in other neuronal types.

As a first step to defining where in the neuron ZDHHC17 might palmitoylate DLK and NMNAT2, we assessed localization of lentivirally expressed HA-ZDHHC17 in cultured DRG neurons. HA-ZDHHC17 was detected in sensory neuron cell bodies, but not axons (Fig 5A). Consistent with this somatic localization, endogenous ZDHHC17 was detected in biochemical fractions from cell body compartments of DRG sensory neuron microfluidic cultures, but was not detected in fractions from distal axon compartments (Fig 5B, C). The axonal proteins tubulin and GAP-43 were readily detected in both soma and distal axon fractions, while the nuclear protein Histone H3 was not present in distal axon fractions, supporting the fidelity of the preparation (Fig 5B). Within the soma, HA-ZDHHC17 colocalized extensively with the Golgi marker GM130 (Fig 5D), consistent with prior studies from non-neuronal cells (Ernst et al. 2018; Ohno et al. 2006). These findings suggest that ZDHHC17 likely palmitoylates DLK and NMNAT2 on somatic Golgi membranes. This conclusion is consistent with prior reports that vesicles upon which wt (i.e. palmitoylation-competent) forms of both DLK and NMNAT2 reside are likely Golgi-derived (Holland et al. 2016; Milde et al. 2013).

**Figure 5:**
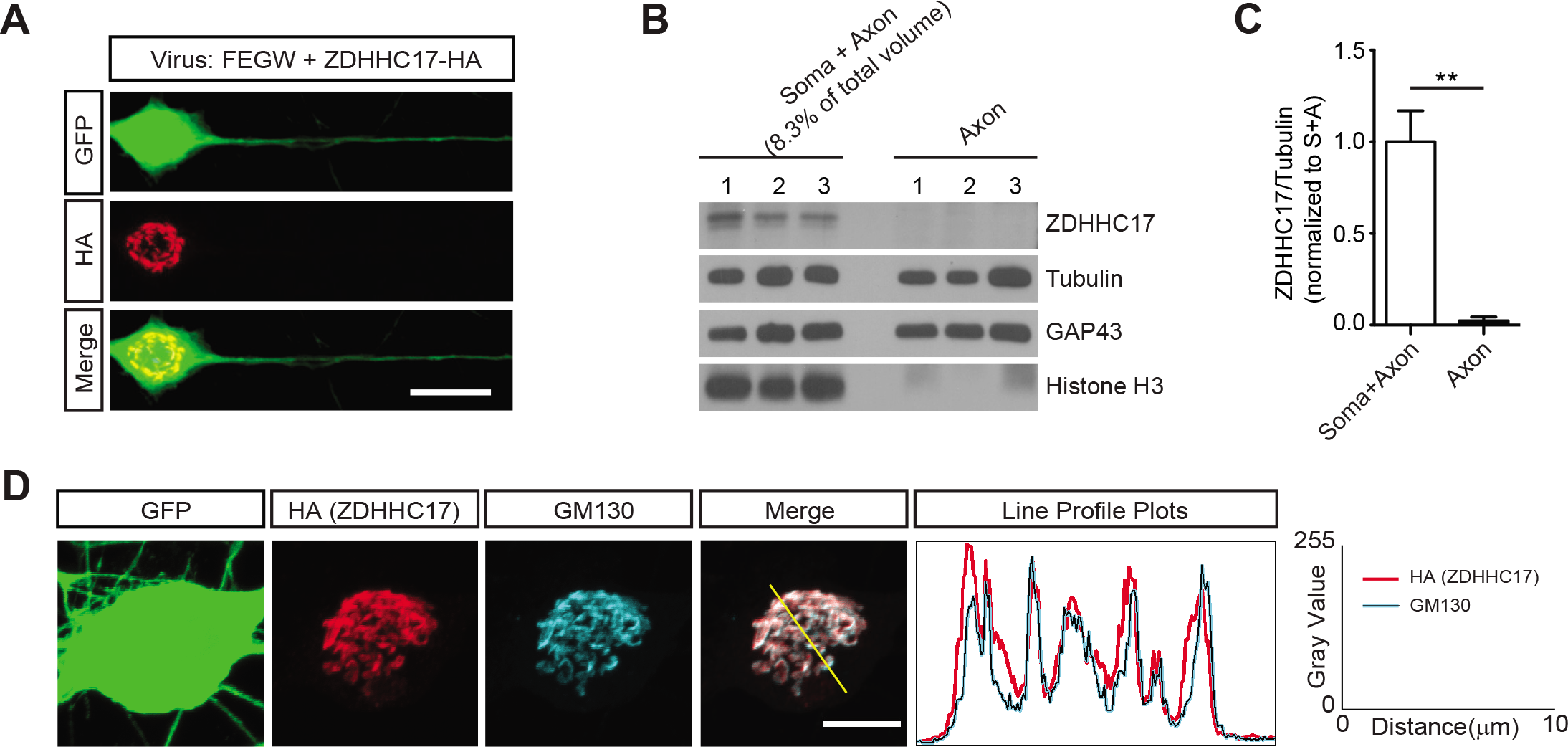
ZDHHC17 Localizes to the Somatic Golgi in Mammalian Sensory Neurons. ***A:*** Images of cultured DRG neurons coinfected with lentiviruses to express GFP and HA-ZDHHC17, immunostained with the indicated antibodies. HA-ZDHHC17 is detected in the neuronal soma but not axons. Images are representative of 10 individual neurons examined. Scale bar: 20 μm. ***B:*** Western blots of somata plus axons (Soma + Axon) or distal axons only (‘Axon’) fractions from 3 sets of DRG neurons cultured in microfluidic chambers, blotted with the indicated antibodies. The ‘Axon’ chamber was lysed in 1/12 of the volume used for the ‘Soma + Axon’ chamber to account for the lower amount of material in the former compartment. ***C:*** Quantified data from n=4 determinations per condition of ZDHHC17 protein level, normalized to tubulin, for each of the indicated subcellular chambers from *B*. **: p<0.01, t-test. ***D:*** DRG sensory neurons infected as in *A* were stained with the indicated antibodies. *Right*: Line profiles across the cell soma (yellow line in merged image) confirm overlapping HA-ZDHHC17 and GM130 signals. Scale bar: 20 μm.

### ZDHHC17 is Also Essential for NMNAT-dependent Distal Axon Integrity

We next addressed the functional role of NMNAT2 palmitoylation by ZDHHC17. Constitutive anterograde supply of NMNAT2 normally prevents distal axon degeneration, and NMNAT2 anterograde vesicular transport requires its palmitoylation (Milde et al. 2013). We therefore asked whether loss of ZDHHC17-dependent NMNAT2 palmitoylation affects distal axon integrity. Consistent with this model, when we extended our lentiviral experiments in DRG neurons beyond the times used for TD experiments (9 days versus 6 days post-infection), we observed distal axon degeneration in *Zdhhc17* ‘knockdown’ cultures (Fig 6A-C). A coinfected cytosolic form of the more stable NMNAT1 (cytoNMNAT1, which can compensate for loss of endogenous NMNAT2 (Sasaki et al. 2006; Sasaki et al. 2009; Yang et al. 2015)) protected axons from the effects of *Zdhhc17* knockdown, supporting the hypothesis that this degeneration was due to loss of NMNAT2 palmitoylation. *Zdhhc17* knockdown axons were also protected by supplementing medium with the NMNAT2 product NAD^+^, a pharmacological treatment that compensates for loss of endogenous NMNAT2 (Sasaki et al. 2006; Wang et al. 2015) (Fig 6B, C). These findings are consistent with the notion that impaired NMNAT2 palmitoylation is a key factor in distal axon degeneration induced by ZDHHC17 loss.

**Figure 6.**
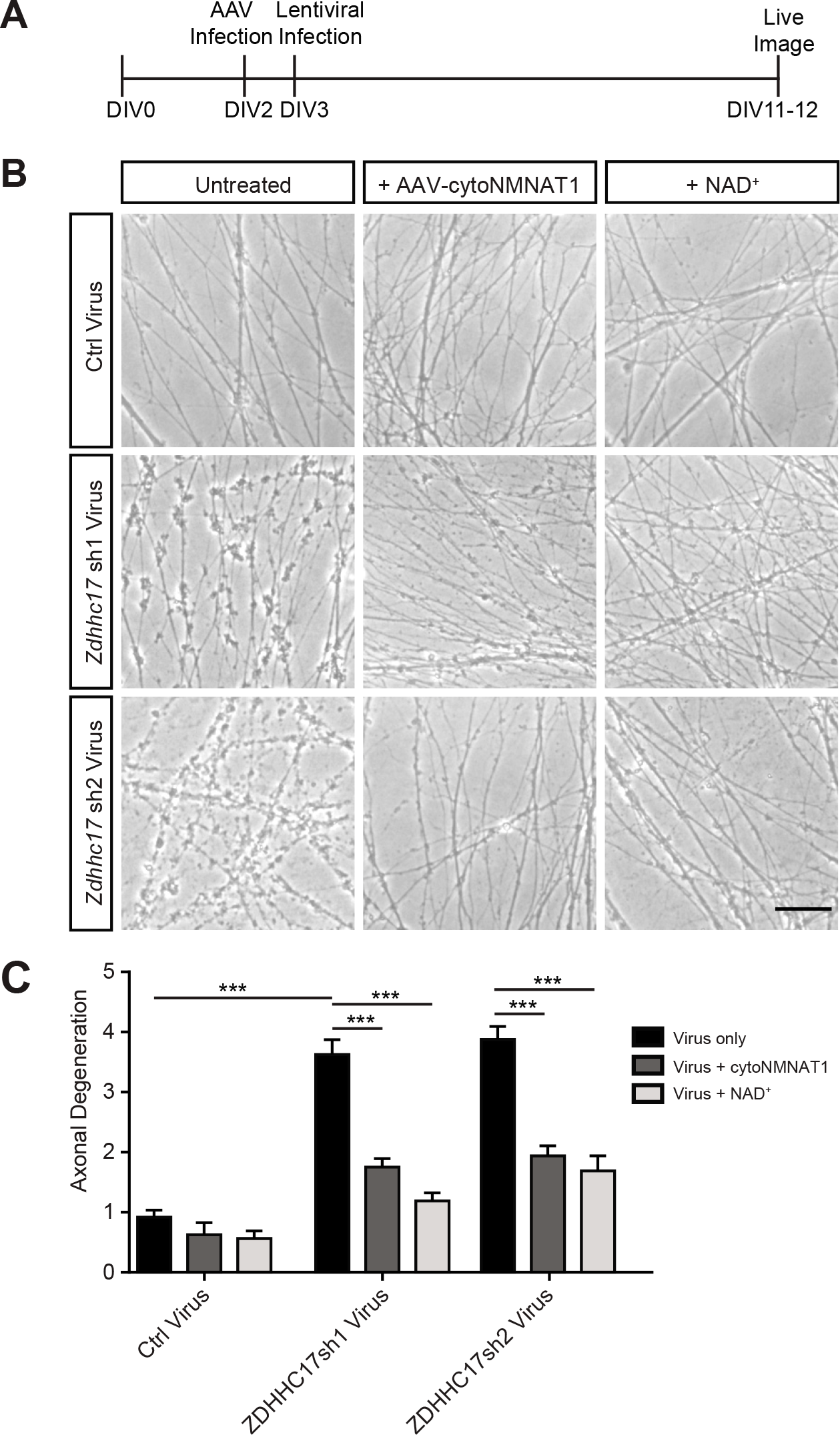
Prolonged *Zdhhc17* Loss Triggers NMNAT-dependent Distal Axon Degeneration in DRG Neurons. ***A:*** Experimental timecourse. Cultured DRG neurons were infected at DIV3 with control lentivirus, or with lentivirus expressing either of two *Zdhhc17* shRNAs. One subset of cultures was pre-infected with AAV expressing HA-tagged cytoNMNAT1. A second subset was fed every 2 days with 1mM NAD+. ***B:*** Example images of distal axons at DIV11-12, from DRG cultures infected with the indicated lentiviruses/AAV or compounds as in *A*. ***C:*** Quantified data from n=3 cultures per condition confirms rescue of axon degeneration in *Zddhc17* knockdown cultures by cyto-NMNAT1 and NAD^+^. ***; p<0.001, n.s.; not significant. One-way ANOVA with Bonferroni *post hoc* test. All data are mean + SEM.

Finally, we asked whether prolonged absence of ZDHHC17 also impacts distal axons *via* loss of palmitoyl-NMNAT2 supply *in vivo*. To address this question, we compared distal optic axon integrity in AAV-GFP- and AAV-Cre-T2A-GFP-injected *Zdhhc17^f/f^* mice. At prolonged times after AAV-Cre-T2A-GFP injection, distal regions of the optic nerve close to the optic chiasm were clearly degenerating, as seen both by fragmentation of GFP-infected axons, and by microgliosis, detected by the marker CD68/ED1 (Ebneter et al. 2010; Palin et al. 2008) (Fig 7A, B). Both the axon degeneration and the microgliosis showed a distal-to-proximal gradient along the optic nerve; distal regions of the nerve close to the optic chiasm were affected, while the proximal region (where the crush occurs in ONC experiments) remained intact until several weeks after *Zdhhc17* CKO (Fig 7B). Neither axon fragmentation nor microgliosis were observed in nerves emanating from control- (GFP-AAV-) infected retinas at any timepoint examined (Fig 7B).

**Figure 7:**
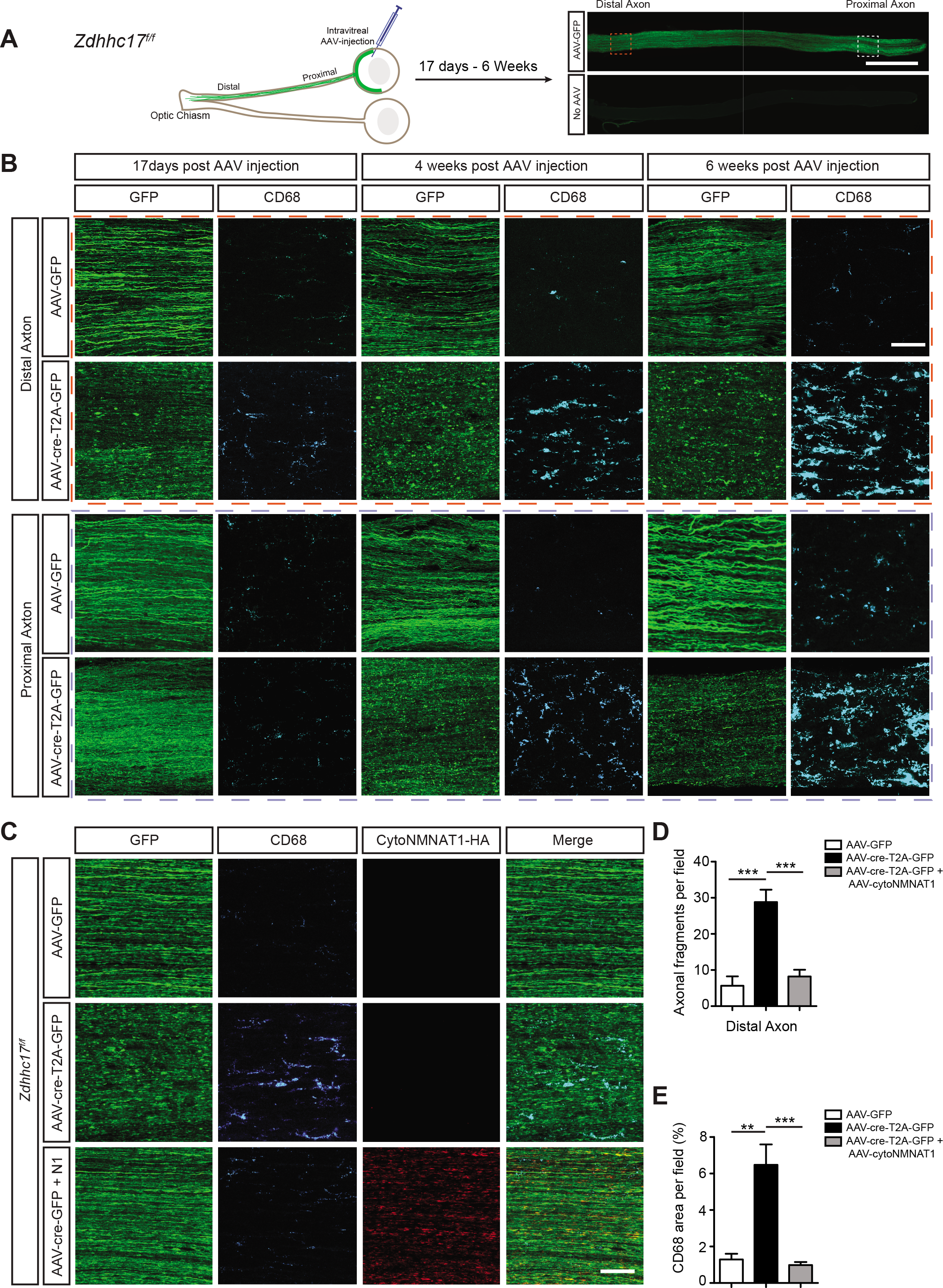
NMNAT-dependent Progressive Distal Axon Degeneration in *Zdhhc17* CKO Optic Nerve. ***A:*** Experimental set-up. *Left: Zdhhc17^f/f^* mice were intravitreally injected with GFP-expressing AAV and optic nerve sections prepared and immunostained. *Right*: Example of optic nerve section at low magnification, from eyes intravitreally injected with AAV-GFP (*top*) or uninjected (*bottom*) and immunostained with anti-GFP antibody. *Orange and purple dashed boxes*: approx. regions used for high magnification images in subsequent panels, respectively. Scale bar: 800 μm. ***B:** Zdhhc17^f/f^* mice were intravitreally injected with the indicated AAVs, fixed at the indicated times and optic nerves were immunostained with the indicated antibodies. AAV-Cre-T2A-GFP, but not AAV-GFP, causes distal-to-proximal axon degeneration, marked by fragmented GFP signal and increase in microgliosis marker CD68. Scale bar: 30 μm. ***C:*** NMNAT-dependent axon degeneration in *Zdhhc17* CKO optic nerve. *Zdhhc17^f/f^* mice were intravitreally injected with the indicated AAVs, fixed at the indicated times and optic nerves were immunostained with antibodies to detect GFP, CD68 and HA-tagged cytoNMNAT1. Scale bar: 30 μm. ***D, E:*** Quantified axon fragmentation and microgliosis in distal optic nerve images from *C*. In mice co-injected with Cre- and HA-cytoNMNAT1 AAVs, axon fragmentation and microgliosis were reduced to control (GFP-AAV-infected) levels.**;p<0.01; ***;p<0.001, one-way ANOVA with *post hoc* Bonferroni test, n=5-8 determinations per condition.

Because the axonal breakdown in *Zdhhc17* CKO optic nerves (Fig 7B) resembled the NMNAT-dependent axon degeneration in cultured DRG neurons (Fig 6B), we asked whether this phenotype could be rescued by genetic manipulation that compensates for loss of NMNAT2. We therefore intravitreally injected AAV expressing Cre-T2A-GFP with or without a second AAV expressing HA-cyto-NMNAT1. Strikingly, HA-cyto-NMNAT1 greatly attenuated both distal axon fragmentation and microgliosis in *Zdhhc17* CKO optic nerves (Fig 7C-E). This result suggests that distal axon degeneration in *Zdhhc17* CKO mice is due to loss of endogenous NMNAT2 palmitoylation.

## Discussion

Previously, separate pathways were thought to control the integrity of RGC somas and distal axons. The prevailing model was that, following optic nerve injury, DLK/JNK retrograde signals triggered somal destruction, while loss of NMNAT2 distal to the injury site activated an intrinsic axon-destruction program. Our findings are consistent with this model but, importantly, reveal that mutual palmitoylation by the same upstream PAT, ZDHHC17, couples the DLK and NMNAT2 signals. ZDHHC17 thus enables palmitoyl-DLK and -NMNAT2 to act in concert, forming an axonal “trust, but verify” system – healthy RGCs send out palmitoyl-NMNAT2 to distal axons to signal that all is well, but simultaneously supply palmitoyl-DLK to detect and respond to axonal insult or injury. This system also ensures a coordinated injury response in cell bodies and distal axons, even though injury/insult-induced physical disruption often prevents communication (by electrical signaling or protein trafficking) between these two subcellular locations.

After injury to the mammalian optic nerve, most RGC cell bodies and axons degenerate, events that our findings suggest depend on ZDHHC17-dependent palmitoylation of DLK and NMNAT2, respectively. This pro-degenerative outcome avoids a situation in which damaged distal axons remain intact in the absence of input, or in which RGCs, which are incapable of long distance regeneration, remain intact but disconnected to their appropriate target(s). Importantly, though, ZDHHC17 also controls palmitoylation and signaling by DLK and NMNAT2 in cultured DRG sensory neurons (Fig. 3, 4, 6) and all three of these proteins are highly expressed in mature DRG neurons *in vivo* (Zeisel et al. 2018). ZDHHC17-dependent regulation thus appears to be equally well-suited to account for the pro-regenerative response to injury that is a hallmark of mature DRG, and other PNS, neurons (Curcio and Bradke 2018; Mahar and Cavalli 2018). Consistent with this notion, injury to the mature mammalian PNS results in NMNAT2-dependent degeneration of severed distal axons (Beirowski et al. 2005), just as in RGCs. However, while the somal response to injury is DLK-dependent in both RGCs and DRG neurons *in vivo*, in the former system DLK upregulates pro-apoptotic mediators (Watkins et al. 2013), while in the latter it upregulates several Regeneration Associated Genes to facilitate axon regeneration (Shin et al. 2012; Shin et al. 2019). ZDHHC17-dependent palmitoylation of DLK and NMNAT2 thus has the potential to ensure appropriate somal and distal axon responses to injury in diverse neuron types, even when the ‘downstream’ functional outcome differs.

### Localization of ZDHHC17 to Somatic Golgi is Critical for the DLK/NMNAT2 System

For the palmitoyl-DLK/NMNAT2 injury/insult response system to function appropriately, the location of the PAT that regulates these two proteins is critical. *A priori*, one might think that this PAT should localize to distal axons, where it could more acutely regulate DLK and/or NMNAT2. However, our assessment of both virally expressed and endogenous ZDHHC17 suggest that this PAT is confined to Golgi-like membranes in the cell soma (Fig 5). This location is highly consistent with observations that wt (i.e. palmitoylation-competent) DLK and NMNAT2 localize to Golgi-derived vesicles (Holland et al. 2016; Milde et al. 2013) and also matches current models of NMNAT2 regulation and function. In particular, NMNAT2 is likely trafficked to distal axons in limiting amounts, with supply constantly balanced by axonal NMNAT2 degradation (Gilley and Coleman 2010; Milde et al. 2013). This ‘limiting supply’ model is consistent with findings that distal axonal regions are most susceptible to degeneration in NMNAT2-dependent dying back axonopathies (Coleman and Freeman 2010), which in turn explains the ‘stockings and gloves’ pathology of these conditions (Argyriou et al. 2014). Significant levels of NMNAT2 PAT (i.e. ZDHHC17) activity in distal axons might facilitate additional NMNAT2 anterograde transport, potentially increasing NMNAT2 levels in extreme distal axons, while reducing NMNAT2 levels in more proximal segments. Such disruption of the axonal NMNAT2 gradient could lead to ‘mixed’ messages regarding the health of proximal and distal axons, an issue circumvented by the somatic Golgi localization of ZDHHC17.

### The ZDHHC17/DLK/NMNAT2 System - a Conserved Regulator of Vertebrate Axon Integrity and Responses to Damage?

The ZDHHC17/DLK/NMNAT2 system is also of interest evolutionarily. ZDHHC17 is highly conserved (Roth et al. 2002; Young et al. 2012) and both DLK and its palmitoyl-site are also evolutionarily ancient (Holland et al. 2016). Our finding that loss of the worm ZDHHC17 orthologs *dhhc-13* and *dhhc-14* markedly affects CeDLK-1 axonal localization (Fig 2), which is itself palmitoylation-dependent (Holland et al. 2016), suggests that ZDHHC17/DLK are a highly evolutionarily conserved enzyme-substrate pair.

In contrast, although NMNATs are also evolutionarily ancient enzymes, the central isoform-specific targeting and interaction domain (ISTID) that distinguishes NMNAT2 is found only in vertebrates (Lau et al. 2010). Interestingly, NMNAT2’s ISTID contains the palmitoyl-sites (C164, C165), the adjacent basic region that is also important for membrane targeting (Milde et al. 2013), and the zDABM that helps confer ZDHHC17-dependent regulation (Lemonidis et al. 2017) (Fig 4B). These three elements are strikingly conserved in all vertebrate NMNAT2 orthologs, but all are absent in invertebrate NMNATs. It thus appears that vertebrates acquired multiple sequence elements that together allow NMNAT2 to propagate ZDHHC17-dependent axon survival signals in vertebrate nervous systems.

### Why is Palmitoylation the Lipid Modification for DLK and NMNAT2 Regulation?

NMNAT2’s ISTID appears well suited to facilitate the vesicle-based anterograde transport and axon survival function of NMNAT2, and vesicle attachment also provides a way for DLK to convey retrograde signals (Holland et al. 2016). Why, though, did palmitoylation arise as the vesicle attachment mechanism for these two proteins, rather than a different lipid modification? One explanation may lie in the observation that palmitate can be added to the central region of a protein sequence, and can thus far more readily regulate protein activity and/or interactions, compared to lipid modifications such as myristoylation and prenylation, which are restricted to the extreme N- and C-termini of proteins, respectively (Montersino and Thomas 2015). In the case of DLK, depalmitoylation causes not just detachment from vesicles but also concomitant loss of kinase activity. Thus, even though palmitoyl-DLK is highly detectable in uninjured neurons, such that palmitoylation is unlikely to be the ‘on’ switch for DLK-JNK signaling *per se* (Holland et al. 2016), the coupling of palmitoylation to DLK kinase activity is a security feature that likely minimizes phosphorylation of inappropriate substrates (Holland et al. 2016; Holland and Thomas 2017; Montersino and Thomas 2015).

Intriguingly, there are suggestions that palmitoylation also controls NMNAT2 activity, but in the opposite way. For example, wt NMNAT2 has a vastly slower (>10-fold) turnover rate (*k_cat_*) than palmitoyl-mutant NMNAT2 in *in vitro* biochemical assays (Mayer et al. 2010) and ISTID-deficient NMNAT2 is also more active than wt NMNAT2 *in vitro* (Lau et al. 2010). These findings suggest that, although palmitoylation is critical for NMNAT2 trafficking on axonal vesicles, NMNAT2 may then need to detach from vesicles in order to increase its catalytic activity and thus maximally protect axons. Indeed, a similar requirement for axonal depalmitoylation was recently suggested to underlie detachment of the microtubule-associated protein MAP6 from vesicles to microtubules (Tortosa et al. 2017). A requirement for NMNAT2 and/or MAP6 detachment from vesicles would exploit a different feature that is unique to palmitoylation, compared to other protein-lipid attachments, namely its reversibility. Identifiying the thioesterase(s) that depalmitolyates NMNAT2 (and DLK) in axons may provide further insights into the control of axonal integrity and is thus an important area for future study.

### Why Might ZDHHC17 be the key PAT for DLK and NMNAT2?

If palmitoylation provides a unique way to couple DLK and NMNAT2 activity to their trafficking, why might a single PAT, and in particular ZDHHC17, regulate these two proteins? One simple explanation is that this arrangement facilitates coordinated regulation of the levels of these three proteins. Consistent with this notion, ZDHHC17 is brain-enriched, matching the nervous system specific expression of DLK and NMNAT2 (Fagerberg et al. 2014; Yue et al. 2014). Moreover, ZDHHC17 expression correlates better with that of both DLK and NMNAT2 than does any other PAT across multiple nervous system cell types (Fig S3C, D and S4E, F). There may, however, be other reasons why ZDHHC17 is the major PAT for DLK and NMNAT2. For example, in addition to DLK and NMNAT2, several other well-characterized ZDHHC17 substrates are axonal and/or pre-synaptic proteins that undergo vesicular transport (Lemonidis et al. 2015). This raises the possibility that ZDHHC17 is enriched within a Golgi subdomain from which such axonal palmitoyl-proteins bud off into vesicles, or through which they pass prior to such a budding step. Imaging studies, most likely at super-resolution, could provide insight into this issue.

### New Roles for ZDHHC17 in Acute Injury Signaling

Our finding that ZDHHC17 is essential for palmitoylation and signaling by both DLK and NMNAT2 advances our understanding of the neuronal roles of this PAT. ZDHHC17 was first identified as an interactor of Huntingtin, the protein whose poly-glutamine expansion underlies Huntington Disease (HD; mutant HTT [mHTT] (Singaraja et al. 2002). The presence of mHTT reduces ZDHHC17 PAT activity and prolonged, global loss of *Zdhhc17* phenocopies several features of HD in mice (Huang et al. 2011; Singaraja et al. 2011). These findings suggest that ZDHHC17 activity is neuroprotective in the context of this chronic neurodegenerative condition, a hypothesis consistent with ZDHHC17’s ability to palmitoylate several key regulators of physiological neuronal function (Huang et al. 2004). However, our data reveal that ZDHHC17 also plays an active, critical role in more acute forms of neurodegeneration. Whether or how ZDHHC17 activity towards pro-survival versus pro-death substrates might be altered in different neuropathological conditions is worthy of further investigation. It is also an intriguing possibility that loss of NMNAT2 palmitoylation (and hence failure to supply palmitoyl-NMNAT2) in peripheral motor axons explains the paralysis and subsequent death of mice in which *Zdhhc17* is knocked out in adulthood (Sanders et al. 2016).

### DLK and NMNAT2 are Dominant ZDHHC17 Substrates Responsible for Somal and Distal Axon Integrity

ZDHHC17 is implicated in other neuropathological situations and, as described above, other ZDHHC17 substrates have been identified (Greaves et al. 2010; Huang et al. 2004). To what extent can we thus attribute the phenotypes that we report to direct regulation of DLK and/or NMNAT2 by ZDHHC17, rather than to other substrates of this PAT? Importantly, the block of retograde axon-to-soma signaling and reduction of DLK palmitoylation are some of the most immediate effects observed following loss of *Zdhhc17*. At this time, cell bodies and axons of *Zdhhc17* knockdown and CKO neurons appear phenotypically normal (J.N., S.S., H-K.J. and G.T., unpublished observations), supporting the hypothesis that direct regulation of the DLK-JNK pathway by ZDHHC17 underlies the block of axon-to-soma signaling that we observe. In contrast, degeneration of distal axons only becomes evident at later times after *Zdhhc17* loss (Fig 6, 7). Other ZDHHC17 substrates may thus contribute to this latter phenotype. Importantly, though, effects of *Zdhhc17* loss on axon integrity are almost completely rescued, both *in vitro* and *in vivo*, by genetic or pharmacological interventions that restore levels of NAD^+^, the key NMNAT2 reaction product (Fig 6, 7). These findings support the hypothesis that NMNAT2 is indeed the key ZDHHC17 substrate whose lack of palmitoylation underlies these more chronic axon degeneration phenotypes.

### A holistic view of the control of neuronal integrity?

Our understanding of processes that govern somal and distal axon degeneration has increased markedly in recent years, with several key molecular players identified (Coleman and Freeman 2010; Gerdts et al. 2016; Wang et al. 2012; Welsbie et al. 2017). Nonetheless, many studies of axonal injury responses in RGCs have assessed only one of these processes in isolation, and on the few occasions when both processes have been addressed, results would initially suggest that they are mediated by different pathways (Fernandes et al. 2014; Fernandes et al. 2018). We propose that the control of somal and distal axon integrity might instead be considered as a single, holistic process, involving two palmitoylation-dependent pathways acting in concert, even when physical injury prevents communication between them. This modified view will be important when considering potential therapeutic interventions for neuropathological conditions that primarily affect distal axons versus neuronal cell bodies.

## Acknowledgments

We thank Drs. Fengsong Qin and Yingpeng Liu for AAV preparation and Dr. C. Su (UCSD) for invaluable discussions. This work was supported by NIH (R01NS094402 and R21EY029386 to G.M.T., R37NS035546 to Y.J. and R01EY024932, R01EY023295 and EY028106 to Y.H.), Shriners Hospitals for Children (Grant 85600 PHI to G.M.T. and a Special Shared Facility Grant for Viral Production to G.M.S.) and BrightFocus Foundation (G2019267, to G.M.T.). S.S.S. acknowledges a Brody Family Medical Trust Fund Fellowship.

## Author Contributions

Conceptualization, Supervision and Funding Acquisition: G.M.S., Y.H., Y.J., G.M.T.

Investigation: J.N., S.S.S., H-K.J., S.M.H., Y.S., K.M.C., L.M.H., G.M.T.

Methodology: H.H., Y.H., G.M.T.

Resources: M.R.H., G.M.S.

## KEY RESOURCES TABLE

| REAGENT or RESOURCE | SOURCE | Cat# | IDENTIFIER |
| --- | --- | --- | --- |
| Antibodies |  |  |  |
| Rabbit anti-phospho c-Jun S63 II | Cell Signaling Technology | 9261 | AB_2130162 |
| Rabbit anti-phospho c-Jun S73 | Cell Signaling Technology | 3270 | AB_2129575 |
| Rabbit anti-phospho c-Jun S63 | Cell Signaling Technology | 91952 |  |
| Rabbit anti-DLK/MAP3K12 | Sigma/ Prestige | HPA039936 | AB_10795239 |
| Rabbit anti-DLK/MAP3K12 | Thermofisher Scientific | PA5-32173 | AB_2549646 |
| Mouse anti-GFP | Life Technologies | A11120 | AB_221568 |
| Rabbit anti-GFP | Life Technologies | A11122 | AB_221569 |
| Mouse anti-Tubulin β3 | BioLegend/Covance | MMS-435P | AB_2313773 |
| Mouse anti-Myc 9E10 | UPenn Cell Center | 3207 |  |
| Mouse anti-Myc 9E10 | Enzo Life Sciences | ENZ-ABS463-0200 |  |
| Rabbit anti-Myc | Cell Signaling Technology | 2278 | AB_490778 |
| Mouse anti-HA11 | BioLegend | 901502 | AB_2565007 |
| Rabbit anti HA | Cell Signaling Technology | 3724 | AB_1549585 |
| Mouse anti-tubulin | Millipore Sigma | T7451 | AB_609894 |
| Sheep anti-NGF | CedarLane | CLMCNET-031 | AB_10060173 |
| Rabbit anti-RBPMS | Millipore Sigma | SAB2101964 | AB_2687403 |
| Rabbit anti-ZDHHC17 | Millipore Sigma | H4714 | AB_1858997 |
| Mouse anti-Brn3a | Millipore Sigma | MAB1585 | AB_94166 |
| Rat anti-CD68 | Bio-Rad | MCA341GA | AB_323909 |
| Rabbit anti-GAP43 | Novus | NB300-143 |  |
| Mouse anti-NMNAT2 (B-10) | Santa Cruz Biotechnology | sc-515206 |  |
| Mouse anti-GM130 | BD Biosciences | 610822 | AB_398141 |
| Mouse anti-histone H3 | Cell Signaling Technology | 4499 | AB_10544537 |
| Rabbit anti-ZDHHC5 | Sigma/ Prestige | HPA014670 | AB_2257442 |
| Rabbit anti-ZDHHC8 | Santa Cruz Biotechnology | sc-86947 | AB_2288678 |
| Bacterial and Virus Strains |  |  |  |
| Biological Samples |  |  |  |
| Chemicals, Peptides, and Recombinant Proteins |  |  |  |
|  | Critical Commercial Assays |  |  |
|  | Deposited Data |  |  |
|  | Experimental Models: Cell Lines |  |  |
| Human: Human Embryonic Kidney (HEK) 293T cells | ATCC | CRL-3216 | CVCL_0063 |
| AAVpro® 293T Cell Line | Takara Bio | 632273 |  |
|  | Experimental Models: Organisms/Strains |  |  |
| Mouse: *Zdhhc17<sup>fl/fl</sup>* in FVB background strain | Sanders *et al.*, 2016 |  |  |
| *C. elegans* strain CZ10969, genotype *mec-7-GFP(muls32) II* |  |  | CZ10969 |
| *C. elegans* strain CZ26963, genotype *mec-7-GFP(muls32) II ; dhhc-13(ju1673) IV* | This study |  | CZ26963 |
| *C. elegans* strain CZ19203, genotype *mec-7-GFP(muls32) II ; dhhc-13(gk36) IV* | This study |  | CZ19203 |
| *C. elegans* strain CZ19204, genotype *mec-7-GFP(muls32) II ; dhhc-14(gk330) X* | This study |  | CZ19204 |
| *C. elegans* strain CZ26287, genotype *mec-7-GFP(muls32) II ; dhhc-13(gk36) IV ; dhhc-14(gk330) X* | This study |  | CZ26287 |
| *C. elegans* strain CZ26964, genotype *mec-7-GFP(muls32) II ; dhhc-13(ju1673) IV ; dhhc-14(gk330) X* | This study |  | CZ26964 |
| *C. elegans* strain CZ20310, genotype *Pmex-5-His-72::miniSOG(juSi164) unc-119(ed3) III* |  |  | CZ20310 |
| *C. elegans* strain CZ23498, genotype *Pmec-4-GFP::DLK-1L Pmec-4-tagRFP(juls501)* |  |  | CZ23498 |
| *C. elegans* strain CZ26965,<br>genotype *dhhc-13(ju1673) IV*;<br>*Pmec-4-GFP::DLK-1L Pmec-4-*<br>*tagRFP (juls501)* | This study |  | CZ26965 |
| *C. elegans* strain CZ26344,<br>genotype *dhhc-13(gk36) IV*;<br>*Pmec-4-GFP::DLK-1L Pmec-4-*<br>*tagRFP (juls501)* | This study |  | CZ26344 |
| *C. elegans* strain CZ26345,<br>genotype *dhhc-14(gk330) X*;<br>*Pmec-4-GFP::DLK-1L Pmec-4-*<br>*tagRFP (juls501)* | This study |  | CZ26345 |
| *C. elegans* strain CZ26346,<br>genotype *dhhc-13(gk36) IV*;<br>*dhhc-14(gk330) X*; *Pmec-4-*<br>*GFP::DLK-1L Pmec-4-*<br>*tagRFP(juls501)* | This study |  | CZ26346 |
| *C. elegans* strain CZ26966,<br>genotype *dhhc-13(ju1673) IV*;<br>*dhhc-14(gk330) X*; *Pmec-4-*<br>*GFP::DLK-1L Pmec-4-*<br>*tagRFP(juls501)* | This study |  | CZ26966 |
| Oligonucleotides |  |  |  |
| Recombinant DNA |  |  |  |
| pAAV-GFP | This study |  |  |
| pAAV-GFP-DLKsh | This study |  |  |
| pAAV-Cre-T2A-GFP | This study |  |  |
| pAAV-wtDLK-HA | This study |  |  |
| pAAV-DLK-CS-HA | This study |  |  |
| pAAV-cytoNMNAT1-HA | This study |  |  |
| pAAV2 (pACG2)-RC triple mutant | Petrs-Silva et al., 2011 |  |  |
| pHelper | Agilent | 240071 |  |
| pFEGW | Holland *et al.*, 2016 |  |  |
| pFEGW DLKsh | Holland *et al.*, 2016 |  |  |
| pFEW wtDLK-myc | Holland *et al.*, 2016 |  |  |
| wtDLK in pEGFP-N1 | Holland *et al.*, 2016 |  |  |
| DLK-CS in pEGFP-N1 | Holland *et al.*, 2016 |  |  |
| DLK-PVAA in pEGFP-N1 | This study |  |  |
| pFEW HA-Zdhhc3 | This study |  |  |
| pFEW HA-Zdhhc5 | This study |  |  |
| pFEW HA-Zdhhc10 | This study |  |  |
| pFEW HA-Zdhhc14 | This study |  |  |
| pFEW HA-Zdhhc15 | This study |  |  |
| pFEW HA-Zdhhc17 | This study |  |  |
| pFEW Zdhhc17-HA | This study |  |  |
| pFEW Zdhhc17-N100A-HA | This study |  |  |
| wtNMNAT2 in pEGFP-N1 | This study |  |  |
| pFEW wtNMNAT2-myc | This study |  |  |
| pFEW NMNAT2-4A-myc | This study |  |  |
| pFEW NMNAT2-3BR-myc | This study |  |  |
| pFEW NMNAT2-4A3BR-myc | This study |  |  |
| Software and Algorithms |  |  |  |
| NIH ImageJ | NIH Image |  | SCR_003070 |
| Fiji | Laboratory for Optical and Computational Instrumentation |  | SCR_002285 |
| GraphPad Prism | GraphPad Software Inc. |  | SCR_002798 |
| Other |  |  |  |

## STAR Methods

### Animals

Wild type C57/B6 and *Zdhhc17^f/f^* mice (latter on FVB background; (Sanders et al. 2016)) were housed in a barrier facility in the Lewis Katz School of Medicine at Temple University. Timed-pregnant female Sprague Dawley rats (Strain code 400, Charles River) used for dissociation of embryonic DRG neurons were sacrificed at E16 as previously described (Holland et al. 2016). All procedures involving animals followed National Institutes of Health guidelines and were approved by the Institutional Animal Care and Use Committee (IACUC) of Temple University.

### AAV Vectors and Preparation

Vector pAAV-hSyn-hChR2(H134R)-eYFP (Addgene plasmid #26973) was modified to replace the hChR2(H134R)-eYFP fragment with shRNA resistant forms of wtDLK or DLK-CS carrying a C-terminal Hemagglutinin (HA) tag. Generation of shRNA-resistant wtDLK and DLK-CS was previously described (Holland et al. 2016) and the HA epitope was added by PCR. The same parent plasmid was also cut with *MluI and HindIII* to replace the hSyn-hChR2(H134R)-eYFP fragment with a cassette (synthesized by Genewiz) containing an H1 promoter and DLK shRNA flanked by *PacI* restriction sites (Holland et al. 2016) followed by the human synapsin promoter, an additional PspXI site, Kozak sequence and eGFP cDNA. The resultant vector, termed pAAV-DLKsh-GFP was cut with *PacI* and re-ligated to remove the H1 promoter / DLKshRNA fragment, generating pAAV-GFP. pAAV-GFP was cut with PspXI / HindIII to replace eGFP with a cassette coding for Cre recombinase followed by a self-cleaving T2A sequence and eGFP (synthesized by Genewiz). The resultant vector was termed pAAV-Cre-T2A-GFP. pAAV-GFP was also cut with PspXI / HindIII to replace eGFP with a cassette coding for a cytosolic form of NMNAT1 (Sasaki et al. 2006) with a C-terminal HA-tag (synthesized by Genewiz).

Adeno-associated viruses were made in HEK293T cells by co-transfecting each of the above plasmids with pAAV2 (pACG2)-RC triple mutant (Y444, 500, 730F) (Petrs-Silva et al. 2011)and pHelper (Stratagene, La Jolla, California) plasmids. Cells were lysed 72h post-transfection to release viral particles, which were precipitated using 40% (w/v) polyethylene glycol and purified by cesium chloride density gradient centrifugation. Fractions with refractive index from 1.370 to 1.374 were dialyzed in MWCO 7000 Slide-A-Lyzer cassettes (Thermo Fisher Scientific, Waltham, Massachusetts) overnight at 4°C. AAV titers used for this study were in the range of 1.5–2.5 x 10^12^ genome copies (GC)/ml determined by real-time PCR.

### Lentiviral Vectors and Preparation

Lentiviral vectors carrying DLK shRNA or shRNA-resistant DLK rescue constructs were previously described (Holland 2016). cDNAs for mouse Zdhhc3, Zdhhc5, Zdhhc7, Zdhhc10, Zdhhc14, Zdhhc15 (all kind gifts of Masaki Fukata (Fukata et al. 2004)) were subcloned from their original vectors into modified FEW vector downstream of an N-terminal HA tag. A previously reported Zdhhc17 cDNA (Fukata et al. 2004) was subcloned into modified FEW vector downstream of a C-terminal HA tag (termed ‘FEW-HA’) with the addition of the N-terminal sequence atgcagcgggaggagggatttaacaccaag (corresponding amino acid sequence: MQREEGFNTK). This sequence includes the first potential Kozak/start site, as previously reported (Huang et al. 2004; Singaraja et al. 2002). Zdhhc17 N100A mutant (Verardi et al. 2017) was generated by Splicing by Overlap Extension (SOE) PCR using the Zdhhc17 cDNA template and the resultant product was subcloned into FEW-HA. DLK-GFP cDNA was previously described (Holland et al. 2016). The DLK-PVAA-GFP mutant was made by replacing the wtDLK-GFP XhoI-EcoRI fragment with a similar fragment (synthesized by Genewiz) in which codons coding for Met134 and Ile135 were both mutated to code for Alanine. The cDNA of wt rat NMNAT2 was synthesized by Genewiz with XhoI and NotI extensions at the 5’ and 3’ end, respectively and subcloned into FEW-myc and FEW-GFP vectors. NMNAT2-4A mutant was made by replacing the wtNMNAT2 XhoI-BstEII fragment with a similar fragment (synthesized by Genewiz) in which codons coding for Pro124, Val125, Gln128 and Pro129 were all mutated to code for Alanine. NMNAT2-3BR mutant was made by replacing the wtNMNAT2 BstEII-NotI fragment with a similar fragment (synthesized by Genewiz) in which codons coding for Lys152, Lys155 and Arg172 were all mutated to code for Alanine (‘3BR fragment). The NMNAT2 4A-3BR mutant was made by removing the BstEII-NotI fragment of NMNAT2-4A and replacing it with the 3BR fragment described above.

One shRNA against rat Zdhhc17 (sequence 5’-ATGAATGCCAGGAGATACAAGCACTTTAA-3’) was purchased from Origene and subcloned into lentiviral vector FEGW (Holland et al. 2016) together with its neighboring U6 promoter. A cassette containing a second shRNA against rat Zdhhc17 (sequence: 5’-CATTAAAGCTACAGAAGAA-3’ plus a neighboring H1 promoter was gene synthesized (Genewiz) and subcloned into FEGW. ShRNA against rat Zdhhc5 in vector FUGW was previously described (Thomas et al. 2012). ShRNA against rat Zdhhc8 (5’-CAGGATGCCACTCTCAGTGAGCCTAAAGC-3’; Origene) was used in its original vector. Lentiviral particles from all cDNA- and shRNA-expressing vectors were generated as previously described (Holland et al. 2016).

### Intravitreal Injection, Optic Nerve Crush and Whole mount Retina Immunostaining

Intravitreal injection of AAV and subsequent optic nerve crush were conducted as previously described (Miao et al. 2016). Briefly, mice were anesthetized with 0.01 mg xylazine + 0.08 mg ketamine per gram of body weight. For studies of c-Jun phosphorylation and RGC viability involving DLK and its palmitoylation (Fig 1, Fig S1), 2 μL AAV was injected into the vitreous chamber of 4 week old C57Bl/6 mice. Optic nerve crush (ONC) was performed 10 days after AAV injection. Three days (for assessment of c-Jun phosphorylation) or 3 weeks (for assessment of RGC viability) post-ONC, mice were again anesthetized with xylazine/ketamine as above and transcardially perfused with 4% paraformaldehyde (PFA) in PBS. For studies in *Zdhhc17^f/f^* mice, 37-day-old mice were anesthetized as above. Optic nerve crush was performed 9 days post-injection and mice were reanesthetized and perfused 33 hours post-ONC.

In all *in vivo* experiments, following perfusion with 4% PFA/1 x PBS, eyeballs were subsequently post-fixed for another 2 h prior to dissection of retinas. Whole mount retinal staining was performed as previously described (Miao et al. 2016). Briefly, wide-field images of flat-mounted retinas were acquired using a Nikon 80i epi-fluorescent microscope with a 10x objective (experiments in Fig 1, Fig 4G and Fig S1) or using a Leica SP8 confocal microscope with a 40x oil immersion objective (experiments in Fig 3G and Fig. 7).

For studies of optic nerve integrity, mice were anesthetized and intravitreally injected with AAV as above. At 17 days, 4 weeks or 6 weeks post-AAV injection, mice were re-anesthetized and transcadially prefused as above. Optic nerves (ONs) were isolated and subjected to postfixation with 4% PFA at 4°C overnight. ONs were then washed with PBS and incubated in 30% sucrose solution (in PBS) for at least 24 hr at 4°C for cryoprotection, and then were embedded in Optimal Cutting Temperature (OCT) medium and frozen on dry ice. Thirty-micron-thick sections were prepared using a cryostat, and collected in PBS. Free-floating ON sections were immunostained with GFP, HA and CD68 antibodies.

### Lentiviral Infection, Trophic Deprivation and Imaging of DRG Neuron cultures

Dorsal root ganglion (DRG) neurons were dissociated from E16 rat embryos and conventional ‘mass’ cultures were prepared as previously described (Holland et al. 2016). For TD experiments, neurons were infected with lentivirus at two days *in vitro* (DIV2) and subjected to Trophic Deprivation (TD) at DIV6 or DIV8 (latter timepoint used for *Zdhhc17* shRNA experiments to ensure complete *Zdhhc17* knockdown). For all TD experiments, NGF-containing medium was replaced with fresh Neurobasal medium lacking NGF but containing B27 supplement plus 25μg/mL sheep anti-NGF antibody. Neurons were subsequently lysed or fixed at the timepoints indicated in individual Figures. For studies of NMNAT2 distribution (Fig. 4), neurons were infected with lentivirus at DIV6 and fixed at DIV8. For studies of HA-ZDHHC17 distribution, neurons were virally infected at DIV4 and imaged at DIV10. For studies of endogenous ZDHHC17 distribution, microfluidic cultures were prepared as previously described (Holland et al. 2016)and the ‘soma + axons’ and ‘distal axons’ chambers were lysed by addition of SDS sample buffer. The ‘Axons’ chamber was lysed in 1/12 of the volume used for the ‘Soma + Axons’ chamber to account for the lower amount of material in the former compartment.

For studies of long-term axonal viability (Fig 6), DRG neurons were lentivirally infected on DIV2. In some experiments, sister cultures were infected with AAV expressing HA-cytoNMNAT1 on DIV1. In some experiments, 1mM NAD+ was added on alternate days after lentiviral infection. In all cases, bright field images were acquired on DIV11-12.

For immunocytochemical readouts, dissociated DRG neurons cultured on coverslips were rinsed once with 1x Recording buffer (25mM HEPES pH7.4, 120mM NaCl, 5mM KCl, 2mM CaCl_2_, 1mM MgCl_2_, 30mM Glucose) and fixed in 4% paraformaldehyde (PFA)/sucrose for 10min at room temperature. Samples were permeabilized in PBS containing 0.25% (w/v) Triton-X-100 for 10 min at 4°C, blocked with PBS containing 10% (v/v) Normal Goat Serum (SouthernBiotech, 0060-01) for 1 hour and incubated in primary antibodies overnight at 4°C in blocking solution. After 3 washes with PBS, cells were incubated for 1 hour at room temperature with AlexaDye-conjugated fluorescent secondary antibodies diluted in blocking solution, prior to 3 final PBS washes and mounting in FluorSave reagent (Millipore Sigma).

### Image Analysis of Retinas and Optic Nerves

To quantify c-Jun phosphorylation in AAV-infected retinas for experiments involving DLK knockdown and rescue, images of GFP and P-c-Jun fluorescent signals were thresholded in Fiji (Schindelin et al. 2012) and GFP/P-c-Jun double-positive cells per field were counted. To quantify surviving RGCs, images of GFP and RBPMS fluorescent signals were thresholded in Fiji and GFP/RBPMS double positive cells per field were counted. To quantify c-Jun phosphorylation in AAV-infected retinas for experiments involving *Zdhhc17* CKO, images were thresholded as above and GFP-positive RGCs were scored as P-c-Jun-positive or −negative by an experimenter blinded to genotype. In each case, 3-4 images were acquired per individual retina and the number of cells per image was averaged to generate a single determination.

Degeneration of optic nerve axons was quantified using optic nerve confocal images from mice that had been intravitreally injected with AAV to express GFP plus/minus Cre. The 5 consecutive most intensely GFP-positive slices for each mouse were marked and each of the five images was then thresholded to an identical value across conditions. Infected (GFP-positive) axonal fragments (‘blebs’) were quantified by analyzing particles of defined size (size: 10-infinity pixels, circularity: 0.40~1.00) in ImageJ for each of the 5 z-slice images per individual sample. The number of axon blebs per field was then averaged to generate a single determination. To quantify microgliosis, CD68 signal was analayzed in the same z-slice images. Each image was again thresholded to an identical value and CD68-positive signal was detected by analyzing particles (size: 5-infinity, circularity: 0.0-1.0) in ImageJ. The CD68-positive area was then plotted as a percentage of the total area of the field imaged.

### Image Analysis in Cultured Neurons

c-Jun phosphorylation in cultured DRG neurons was quantified using ImageJ’s cell batch count plugin. Images were thresholded for phospho c-Jun and NeuN signal and the percentage of phospho c-Jun positive cells that were also NeuN-positive was calculated. To quantify NMNAT2 axonal distribution, maximum intensity projections of confocal images of NMNAT2wt-myc or NMNAT2-4A-3BR-myc were thresholded to an identical value in ImageJ. Tuj1 signal from the same images was separately thresholded (to a different absolute value, but again identically across conditions). Regions of the myc and Tuj1 images that contained cell somas were manually identified and cropped. The thresholded Tuj1 signal was used as a mask and the number of NMNAT2-myc puncta that overlapped with the Tuj1 mask was counted by analyzing particles (size: 6-30 pixels, circularity: 0.6-1.0) in ImageJ.

To assess HA-ZDHHC17 distribution, maximum intensity projections of confocal images of were generated and line intensity profile plots from HA and GM130 (Golgi marker) channels were overlaid in ImageJ.

To quantify axon degeneration, images of distal axons were acquired live and subsequently ranked on a 5-point scale (0 = no degeneration to 5 = severe degeneration), according to the degree of observed axon fragmentation and beading, similar to a prior report (Yang et al. 2015).

### Detection of Palmitoylation by Acyl Biotinyl Exchange (ABE) assay

ABE assays and subsequent western blotting were performed as previously described (Holland, 2016).

### HEK293T Cell ABE and Co-immunoprecipitation Experiments

HEK293T cells were transfected using a calcium phosphate based method as described (Holland et al. 2016). Cells were processed for ABE assays as described (Holland et al. 2016). Co-immunoprecipitations were also performed similar to (Holland et al. 2016). Briefly, HEK293T cells were transfected with C-terminally GFP-tagged DLK cDNA plus the indicated HA-tagged ZDHHC cDNAs or empty vector. All ZDHHC cDNAs were N-terminally tagged except for ZDHHC17, which was C-terminally tagged. The following day cells were lysed with ice-cold Immunoprecipitation buffer (IPB; 20 mM Tris pH 7.5, 150 mM NaCl 0.1% Triton X-100 plus protease and phosphatase inhibitors). Lysates were incubated at 4 degrees for 30 min while rotating prior to centrifugation at 13,000 x *g* and passage through a SpinX column to remove debris. After centrifugation of lysates, a fraction of each supernatant were diluted in sample buffer, denatured, and run on PAGE gels as ‘inputs.’ Remaining supernatants were immunoprecipitated for 90 minutes at 4°C with anti-HA antibodies that had been precoupled to protein G Sepharose beads (GE Healthcare). Beads were washed three times with ice cold IPB containing 0.25M NaCl and twice with IPB. Proteins were eluted with SDS sample buffer and subjected to SDS-PAGE and subsequent immunoblotting.

### HEK293T Cell Immunocytochemical Experiments

Distribution of DLK-GFP in HEK293T cells was assessed by fixing and permeabilizing cells on coverslips 8h after calcium phosphate-based transfection as above. Incubation with primary and secondary antibodies was performed as described for cultured neurons above.

### Assessment of Co-expression of DLK and NMNAT2 with ZDHHC-PATs

Expression of *DLK(Map3k12), Nmnat2* and all 23 mouse *Zdhhc*-PATs from 265 identified nervous system cell subtypes was extracted from mousebrain.org (an online resource containing single cell RNA-Seq data from (Zeisel et al. 2018) and related studies). *Map3k12* expression values in each of the 265 cell types were then plotted in GraphPad Prism against expression values for each PAT individually, generating 23 separate scatter plots. The r-squared value for correlation of each PAT’s expression versus that of *Map3k12* was then calculated in GraphPad Prism and logged. The same analysis was repeated, this time comparing *Nmnat2* expression values versus expression values for each PAT

### Antibodies

Primary antibodies used were as follows: Rabbit anti-phospho c-Jun S63 II (Cell Signaling Technology, #9261, 1:100, used for sensory neuron experiments), Rabbit anti-phospho c-Jun Ser-73 (Cell Signaling Technology, #3270, used for RGC experiments); rabbit anti-DLK/MAP3K12 (Sigma/ Prestige, #HPA039936, used for western blotting); rabbit anti-DLK/MAP3K12 (Thermofisher Scientific, #PA5-32173, used for immunostaining); mouse anti-GFP (Life Technologies, #A11120, clone 3E6 IgG2a, 1:250), rabbit anti-GFP (Life Technologies, #A11122), mouse anti-Tubulin β3 (BioLegend, TUJ1, Ig2a, Covance catalog# MMS-435P, 1:1000), mouse anti-Myc 9E10 (University of Pennsylvania Cell Center, Catalog #3207), mouse anti-HA11 (BioLegend, 901502, IgG1, 1:500), rabbit anti-HA (Cell Signaling Technology, #3724), rabbit anti-myc (Cell Signaling Technology, #2278), mouse anti-tubulin (Millipore Sigma, Catalog #T7451), sheep anti-NGF (CedarLane, catalog #CLMCNET-031). Rabbit anti-RBPMS (Millipore Sigma, #SAB2101964). Rabbit anti-phosphoMKK4 (Cell Signaling Technology, #4514, Rabbit anti-MKK4 (Cell Signaling Technology, #9152), Rabbit anti-phosphoAkt T308 (Cell Signaling Technology, #9275, Rabbit anti-Akt (Cell Signaling Technology, #4691), Rabbit anti-phosphoERK (Cell Signaling Technology, #9106), mouse anti-panERK1/2 (Cell Signaling Technology, #9106), mouse anti-phosphoJNK (Cell Signaling Technology, #9255), rabbit anti pan-JNK (Santa Cruz Biotechnology, #SC-571), rabbit anti-phospho-MAPKAPK2 T334 (Cell Signaling Technology, #3041), rabbit anti-phospho-p38 MAPK Cell Signaling Technology, #4511), rabbit anti-ZDHHC17 (Millipore Sigma #H4714), sheep anti-NGF (Cedarlane #CLMCNET-031), Brn3a (Millipore Sigma #MAB1585), rat anti-CD68, clone ED1 (Bio-Rad #MCA341GA).

### *C. elegans* genetics

*C. elegans* strains were maintained at 20°C on NGM plates as described (Brenner 1974). Pmec-4-GFP::CeDLK-1L(wt)-3’UTR(let-858) (pCZGY2335) was previously described (Holland et al., 2016). The integrated line juIs501 was generated in CZ20310 *Pmex-5-His-72::miniSOG(juSi164) unc-119(ed3) III* background, following the protocol described in (Noma and Jin 2018). Strains are listed in STAR Methods table. Sequences of primers used to genotype individual strains are available upon request.

### Generation of transgenic *C. elegans*

Pmec-4-GFP::CeDLK-1L(wt)-3’UTR(let-858)(pCZGY2335) was previously described (Holland et al., 2016). Multi copy transgenic animals were generated as described (Mello et al. 1991), except that x CZ20310 *Pmex-5-His-72::miniSOG(juSi164) unc-119(ed3) III* was used as host. Pmec-4-GFP::CeDLK-1(wt)-3’UTR(let-858)(pCZGY2335) at 10 ng/ul and Pmec-4-tagRFP (pCZGY2333) at 5 ng/ul were injected into nongravid young adult worms with a coinjection marker Pttx-3-RFP at 85 ng/ul. 12h and 24h after injection, P0 worms were treated with blue LED light for 30 min at 4Hz with 65% power (2mW/mm^2^) (2 light treatments in total). The integrated line juIs501 was isolated by examination of Pttx-3-RFP expression as well as Pmec-4-GFP::DLK-1 and examined for 100% transmission rate in the next generation.

### CRISPR/Cas9 Knock-out of *dhhc-13*

*dhhc-13(ju1673*) was generated following a purified Cas9/crRNA injection protocol. Briefly, sgRNA targeting exon 1 and exon 11 of *dhhc-13* was designed and crRNAs (YJ12386 and YJ12387, IDT Alt-R^®^ CRISPR-Cas9 crRNA, 10μM) were co-injected with *dpy-10* crRNA (5μM), tracrRNA (15μM) and Cas9 protein (28μM). Dpy+ progeny were selected and genotyped for 3.7kb deletion in *dhhc-13* in the next generation (primer sequences available on request). *dhhc-13(ju1673*) was a 3.7kb deletion from exon 1 to exon 10, removing the majority of protein-coding region and is thus considered to be null for *dhhc-13*.

### Fluorescence Microscopy in *C. elegans*

To quantify the subcellular localization of *C. elegans* DLK-1 in PLM touch receptor neuron using *Pmec-4-GFP::DLK-1L Pmec-4-tagRFP(juIs501*), L4 animals were mounted in M9 buffer on 5% agarose pad. GFP-DLK-1L pattern in PLM was scored under a Zeiss Axioplan 2 microscope equipped with Chroma HQ filters and a 63x objective. Visibly detectable GFP puncta in both soma or axon of PLM were scored to define punctate GFP-DLK-1. Diffuse GFP-DLK-1 generally showed decreased fluorescence intensity in axon and soma. All strains were quantified in a genotype-blind manner and data were collected from a minimum of 3 independent observations (30-70 animals per observation) on different days. Example fluorescent images of PLM neurons in Figure S6C were acquired from day 1 adults using a Zeiss LSM710 confocal microscope equipped with a 63x objective. Images are maximal intensity projections from z-stacks (0.5-1 μm/section).

## Supplementary Figure Legends

**Figure S1.**
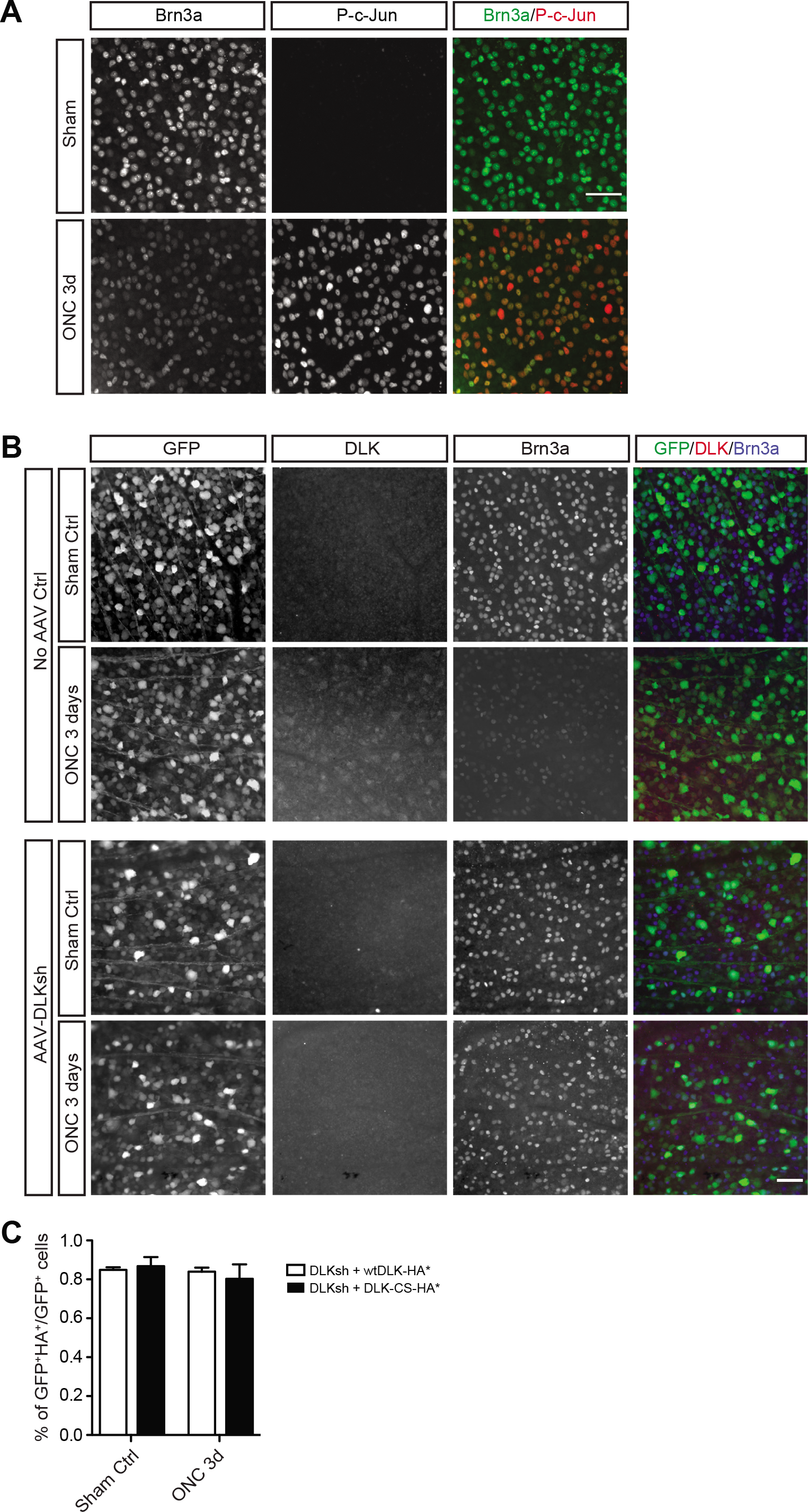
related to Figure 1: Characterization of Optic Nerve Crush (ONC) Response and Confirmation of AAV-mediated DLK knockdown and AAV co-infectivity *in vivo*. ***A:*** Flat-mount retinas were isolated from uninfected mice three days after sham injury (Sham) or optic nerve crush (ONC) and immunostained with the indicated antibodies. ONC markedly downregulates Brn3a, a marker of healthy RGCs, and increases phosphorylation of the injury response transcription factor c-Jun (P-c-Jun), consistent with prior reports (Watkins et al. 2013; Welsbie et al. 2013). Scale bar: 50 μm. ***B:*** Retinas of mice injected with control AAV or AAV-DLKsh were subjected to sham injury or ONC 10 days later. Three days after ONC or sham injury, retinas were isolated and immunostained with the indicated antibodies. DLK is upregulated after ONC in retinas infected with control AAV, consistent with prior reports (Watkins et al. 2013; Welsbie et al. 2013). In contrast, anti-DLK antibody detects only background signals and Brn3a signal is not reduced following ONCin DLK ‘knockdown’ retinas, consistent with (Watkins et al. 2013; Welsbie et al. 2013). Scale bar: 50 μm. ***C:*** Mice were injected with the indicated AAVs and subjected to sham injury or ONC 10 days later as in Fig 1B. Three days later, retinas were isolated and immunostained to detect GFP (indicator of infection with AAV-DLKsh) or HA-tagged DLKwt* or −CS*. Quantified data (4 images per retina, n=3-5 retinas per condition) confirm that >80% of GFP-positive RGCs are also HA-positive i.e. coinfected with both AAVs. All data are mean ± SEM.

**Figure S2.**
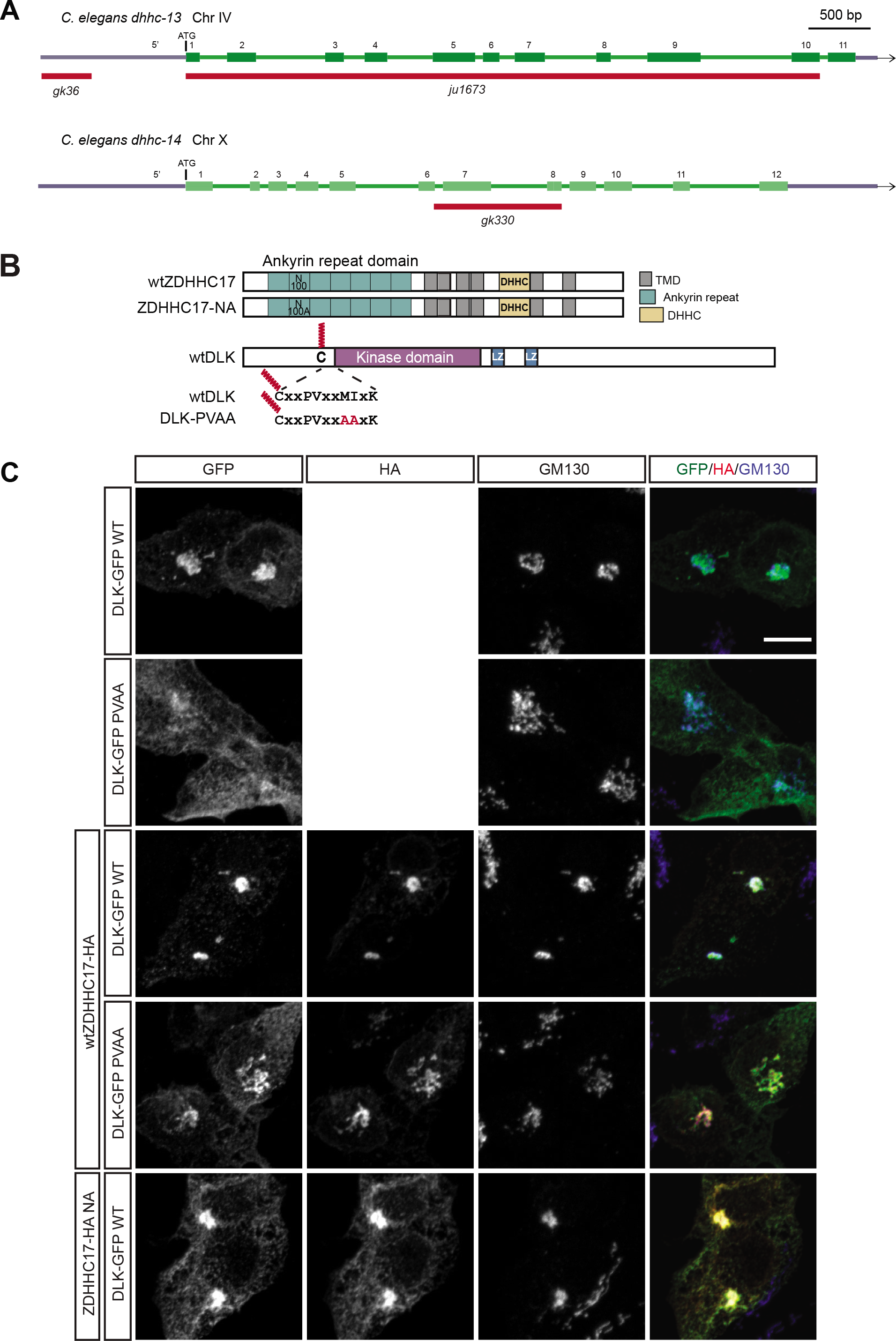
related to Figure 2: Genomic details of *C. elegans* PAT mutant strains and ZDHHC17-dependent distribution of DLK-GFP in HEK293T cells. ***A:*** Schematics of *dhhc-13* and *dhhc-14* genomic loci, with red lines below indicating the locations of mutations. *dhhc-13(gk36*) deletion primarily removes sequences 5’ to ATG. To obtain a definitive null of *dhhc-13*, we generated *dhhc-13(ju1673*) by CRISPR/Cas9, which deletes 3.7kb sequence from exon 1 to exon 10. *dhhc-14(gk330*) deletion removes exons 7 and 8 and is predicted to induce a frameshift, likely leading to genetic null. ***B:*** Schematic of ZDHHC17 and DLK domain structure (reproduced from Fig. 2E), showing locations of mutations disrupting AnkR domain (N100A) and predicted zDABM (PVAA) in ZDHHC17 and DLK, respectively. ***C:*** HEK293T cells transfected to express the indicated cDNAs were immunostained with the indicated antibodies. Right column shows merged images of columns 1-3 for each condition. In HEK293T cells, targeting of DLK-GFP to the Golgi (detected by GM130 marker) is palmitoylation-dependent (Martin et al. 2019). Golgi targeting of DLK-GFP is enhanced by HA-ZDHHC17wt but not by HA-ZDHHC17-N100A mutant, and is also disrupted by DLK zDABM mutation. Scale bar: 10 μm.

**Figure S3.**
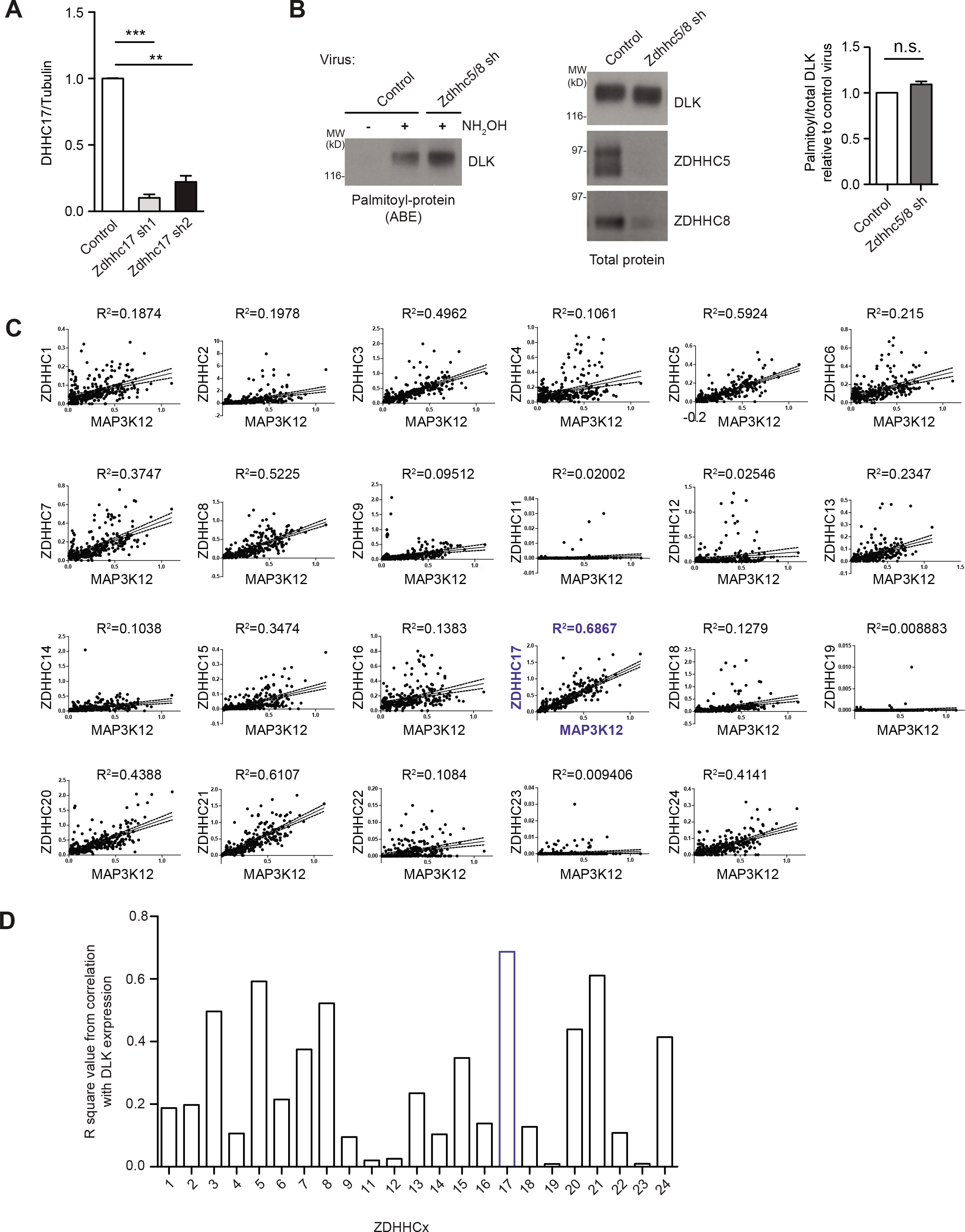
related to Figure 3. Additional Evidence that ZDHHC17 is a Major DLK PAT. ***A:*** Quantified intensities of ZDHHC17 protein levels from Fig 3A, normalized to tubulin, confirm efficacy of *Zdhhc17* shRNAs. N=9-10 individual cultures per condition; ***:p<0.001, **:p<0.01 versus control virus condition, Kruskal Wallis non-parametric test with Dunn’s multiple comparison *post hoc* analysis. ***B:*** Combined knockdown of *Zdhhc5* and *Zdhhc8* does not reduce DLK palmitoylation in DRG neurons. Cultured DRG neurons were infected with control lentivirus (expressing GFP alone) or with lentiviruses expressing GFP plus shRNAs against *Zdhhc5* and *Zdhhc8*. Cultures were lysed 7 days later and processed for ABE. Western blots of ABE fractions (*left*) and total lysates (*right*) reveal that combined *Zdhhc5/8* knockdown does not reduce DLK palmitoylation, despite markedly reduced protein levels of both ZDHHC5 and ZDHHC8. *Right-hand histogram*: quantified data, n=3 determinations per condition. n.s.; not significant, t-test with Welch’s correction. ***C:*** Correlation of *Zddhc17* and DLK (*Map3k12*) expression in the nervous system. DLK expression in 265 mouse nervous system cell types was plotted individually against expression of each of the 23 mouse ZDHHC PATs (from quantitative single cell RT-PCR data; www.mousebrain.org). Linear regression analysis was performed and the r^2^ value was determined for each of the 23 pair-wise comparisons. ***D:*** Histogram of R-squared value for each of the 23 pairwise comparisons in *C* confirms that *Map3k12* expression correlates better with *Zdhhc17* expression than with any other PAT.

**Figure S4.**
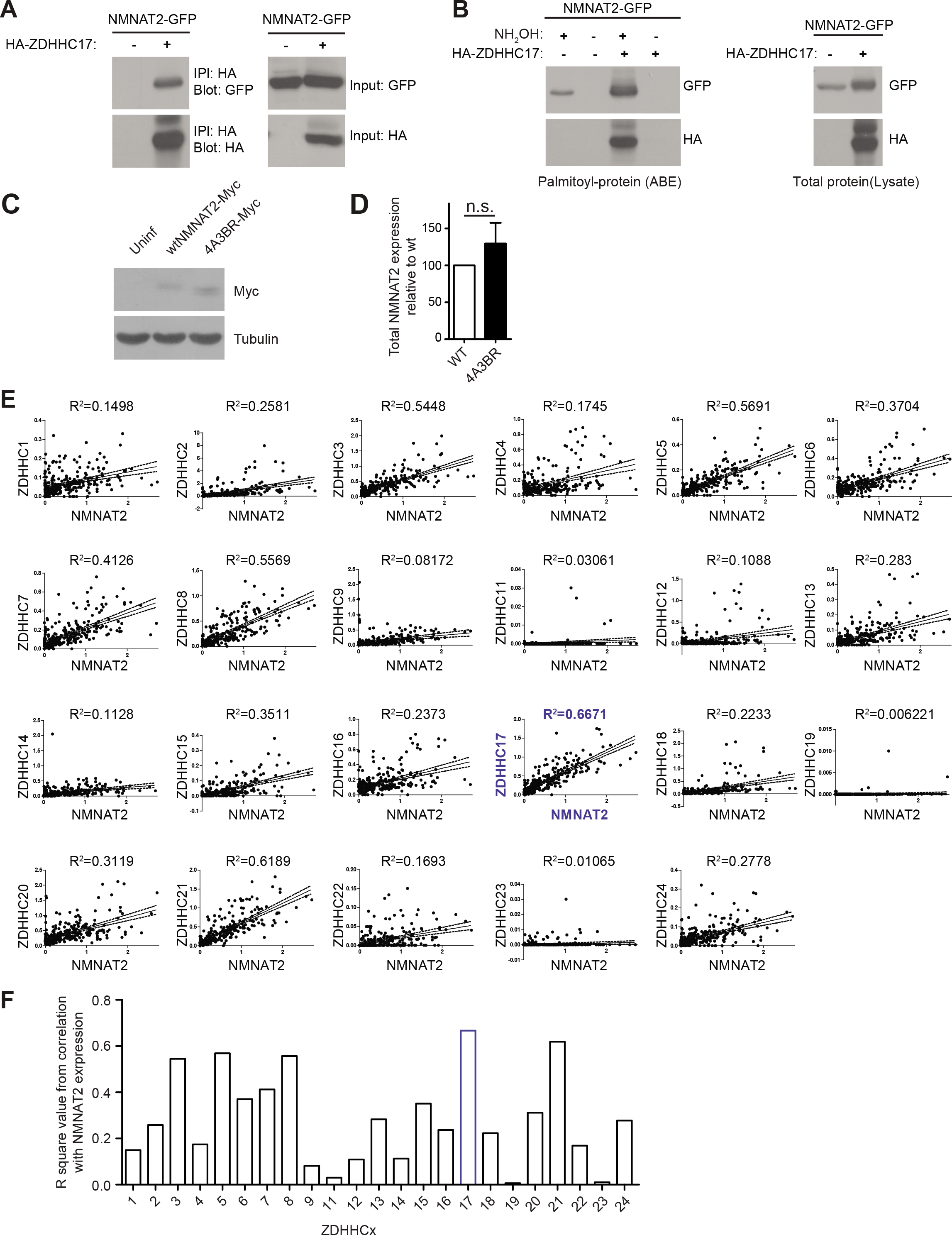
related to Figure 4. Additional Evidence that ZDHHC17 is a Major NMNAT2 PAT. ***A:*** NMNAT2 binds ZDHHC17. Western blots of HA immunopreciptates (*left panels*) and parental cell lysates (*right panels*) from HEK293T cells transfected with the indicated cDNAs and blotted with the indicated antibodies. ***B:*** ZDHHC17 palmitoylates NMNAT2. ABE samples (*left panels*) and total lysates (*right panels*) from HEK293T cells transfected with the indicated cDNAs. Palmitoylation of NMNAT2 is greatly increased by HA-ZDHHC17. ***C:*** Lysates from DRG cultures infected as in Fig 4E were blotted with the indicated antibodies. ***D:*** Quantified NMNAT2-myc expression from n=4 infections per condition from *C* confirms that wt and 4A-3Br forms of NMNAT2-myc express similarly: n.s.: not significant, t-test. ***E:*** Correlation of *Zddhc17* and *Nmnat2* expression in the nervous system. *Nmnat2* expression in 265 mouse nervous system cell types was plotted individually against expression of each of the 23 mouse ZDHHC PATs (from quantitative single cell RT-PCR data, www.mousebrain.org). Linear regression analysis was performed and the r^2^ value was determined for each of the 23 pair-wise comparisons. ***D:*** Histogram of R-squared value for each of the 23 pairwise comparisons in *C* confirms that *Nmnat2* expression correlates better with *Zdhhc17* expression than with any other PAT.

## References

Aguayo AJ, Rasminsky M, Bray GM, Carbonetto S, McKerracher L, Villegas-Perez MP, Vidal-Sanz M, Carter DA. 1991. Degenerative and regenerative responses of injured neurons in the central nervous system of adult mammals. Philos Trans R Soc Lond B Biol Sci 331(1261):337–343.

Argyriou AA, Kyritsis AP, Makatsoris T, Kalofonos HP. 2014. Chemotherapy-induced peripheral neuropathy in adults: a comprehensive update of the literature. Cancer Manag Res 6:135–147.

Beirowski B, Adalbert R, Wagner D, Grumme DS, Addicks K, Ribchester RR, Coleman MP. 2005. The progressive nature of Wallerian degeneration in wild-type and slow Wallerian degeneration (WldS) nerves. BMC Neurosci 6:6.

Beirowski B, Babetto E, Coleman MP, Martin KR. 2008. The WldS gene delays axonal but not somatic degeneration in a rat glaucoma model. Eur J Neurosci 28(6):1166–1179.

Benowitz LI, He Z, Goldberg JL. 2017. Reaching the brain: Advances in optic nerve regeneration. Exp Neurol 287(Pt 3):365–373.

Berger F, Lau C, Dahlmann M, Ziegler M. 2005. Subcellular compartmentation and differential catalytic properties of the three human nicotinamide mononucleotide adenylyltransferase isoforms. J Biol Chem 280(43):36334–36341.

Berkelaar M, Clarke DB, Wang YC, Bray GM, Aguayo AJ. 1994. Axotomy results in delayed death and apoptosis of retinal ganglion cells in adult rats. J Neurosci 14(7):4368–4374.

Blanc M, David F, Abrami L, Migliozzi D, Armand F, Burgi J, van der Goot FG. 2015. SwissPalm: Protein Palmitoylation database. F1000Res 4:261.

Coleman MP, Freeman MR. 2010. Wallerian degeneration, wld(s), and nmnat. Annu Rev Neurosci 33:245–267.

Conforti L, Gilley J, Coleman MP. 2014. Wallerian degeneration: an emerging axon death pathway linking injury and disease. Nat Rev Neurosci 15(6):394–409.

Curcio M, Bradke F. 2018. Axon Regeneration in the Central Nervous System: Facing the Challenges from the Inside. Annu Rev Cell Dev Biol 34:495–521.

Danesh-Meyer HV. 2011. Neuroprotection in glaucoma: recent and future directions. Curr Opin Ophthalmol 22(2):78–86.

Ebneter A, Casson RJ, Wood JP, Chidlow G. 2010. Microglial activation in the visual pathway in experimental glaucoma: spatiotemporal characterization and correlation with axonal injury. Invest Ophthalmol Vis Sci 51(12):6448–6460.

Edmonds MJ, Morgan A. 2014. A systematic analysis of protein palmitoylation in Caenorhabditis elegans. BMC Genomics 15:841.

Ernst AM, Syed SA, Zaki O, Bottanelli F, Zheng H, Hacke M, Xi Z, Rivera-Molina F, Graham M, Rebane AA, Bjorkholm P, Baddeley D, Toomre D, Pincet F, Rothman JE. 2018. S- Palmitoylation Sorts Membrane Cargo for Anterograde Transport in the Golgi. Dev Cell 47(4):479–493 e477.

Fagerberg L, Hallstrom BM, Oksvold P, Kampf C, Djureinovic D, Odeberg J, Habuka M, Tahmasebpoor S, Danielsson A, Edlund K, Asplund A, Sjostedt E, Lundberg E, Szigyarto CA, Skogs M, Takanen JO, Berling H, Tegel H, Mulder J, Nilsson P, Schwenk JM, Lindskog C, Danielsson F, Mardinoglu A, Sivertsson A, von Feilitzen K, Forsberg M, Zwahlen M, Olsson I, Navani S, Huss M, Nielsen J, Ponten F, Uhlen M. 2014. Analysis of the human tissue-specific expression by genome-wide integration of transcriptomics and antibody-based proteomics. Mol Cell Proteomics 13(2):397–406.

Fernandes KA, Harder JM, Fornarola LB, Freeman RS, Clark AF, Pang IH, John SW, Libby RT. 2012. JNK2 and JNK3 are major regulators of axonal injury-induced retinal ganglion cell death. Neurobiol Dis 46(2):393–401.

Fernandes KA, Harder JM, John SW, Shrager P, Libby RT. 2014. DLK-dependent signaling is important for somal but not axonal degeneration of retinal ganglion cells following axonal injury. Neurobiol Dis 69:108–116.

Fernandes KA, Mitchell KL, Patel A, Marola OJ, Shrager P, Zack DJ, Libby RT, Welsbie DS. 2018. Role of SARM1 and DR6 in retinal ganglion cell axonal and somal degeneration following axonal injury. Exp Eye Res 171:54–61.

Gerdts J, Summers DW, Milbrandt J, DiAntonio A. 2016. Axon Self-Destruction: New Links among SARM1, MAPKs, and NAD+ Metabolism. Neuron 89(3):449–460.

Ghosh AS, Wang B, Pozniak CD, Chen M, Watts RJ, Lewcock JW. 2011. DLK induces developmental neuronal degeneration via selective regulation of proapoptotic JNK activity. J Cell Biol 194(5):751–764.

Gilley J, Coleman MP. 2010. Endogenous Nmnat2 is an essential survival factor for maintenance of healthy axons. PLoS Biol 8(1):e1000300.

Gilley J, Orsomando G, Nascimento-Ferreira I, Coleman MP. 2015. Absence of SARM1 rescues development and survival of NMNAT2-deficient axons. Cell Rep 10(12):1974–1981.

Gilley J, Ribchester RR, Coleman MP. 2017. Sarm1 Deletion, but Not Wld(S), Confers Lifelong Rescue in a Mouse Model of Severe Axonopathy. Cell Rep 21(1):10–16.

Greaves J, Gorleku OA, Salaun C, Chamberlain LH. 2010. Palmitoylation of the SNAP25 protein family: specificity and regulation by DHHC palmitoyl transferases. J Biol Chem 285(32):24629–24638.

Harvey AR. 2007. Combined therapies in the treatment of neurotrauma: polymers, bridges and gene therapy in visual system repair. Neurodegener Dis 4(4):300–305.

He M, Abdi KM, Bennett V. 2014. Ankyrin-G palmitoylation and betaII-spectrin binding to phosphoinositide lipids drive lateral membrane assembly. J Cell Biol 206(2):273–288.

Holland SM, Collura KM, Ketschek A, Noma K, Ferguson TA, Jin Y, Gallo G, Thomas GM. 2016. Palmitoylation controls DLK localization, interactions and activity to ensure effective axonal injury signaling. Proc Natl Acad Sci U S A 113(3):763–768.

Holland SM, Thomas GM. 2017. Roles of palmitoylation in axon growth, degeneration and regeneration. J Neurosci Res 95(8):1528–1539.

Hu Y, Park KK, Yang L, Wei X, Yang Q, Cho KS, Thielen P, Lee AH, Cartoni R, Glimcher LH, Chen DF, He Z. 2012. Differential effects of unfolded protein response pathways on axon injury-induced death of retinal ganglion cells. Neuron 73(3):445–452.

Huang K, Sanders SS, Kang R, Carroll JB, Sutton L, Wan J, Singaraja R, Young FB, Liu L, El- Husseini A, Davis NG, Hayden MR. 2011. Wild-type HTT modulates the enzymatic activity of the neuronal palmitoyl transferase HIP14. Hum Mol Genet 20(17):3356–3365.

Huang K, Yanai A, Kang R, Arstikaitis P, Singaraja RR, Metzler M, Mullard A, Haigh B, Gauthier-Campbell C, Gutekunst CA, Hayden MR, El-Husseini A. 2004. Huntingtin-interacting protein HIP14 is a palmitoyl transferase involved in palmitoylation and trafficking of multiple neuronal proteins. Neuron 44(6):977–986.

Huntwork-Rodriguez S, Wang B, Watkins T, Ghosh AS, Pozniak CD, Bustos D, Newton K, Kirkpatrick DS, Lewcock JW. 2013. JNK-mediated phosphorylation of DLK suppresses its ubiquitination to promote neuronal apoptosis. J Cell Biol 202(5):747–763.

Kalesnykas G, Oglesby EN, Zack DJ, Cone FE, Steinhart MR, Tian J, Pease ME, Quigley HA. 2012. Retinal ganglion cell morphology after optic nerve crush and experimental glaucoma. Invest Ophthalmol Vis Sci 53(7):3847–3857.

Kanaani J, Diacovo MJ, El-Husseini Ael D, Bredt DS, Baekkeskov S. 2004. Palmitoylation controls trafficking of GAD65 from Golgi membranes to axon-specific endosomes and a Rab5a-dependent pathway to presynaptic clusters. J Cell Sci 117(Pt 10):2001–2013.

Kwong JM, Quan A, Kyung H, Piri N, Caprioli J. 2011. Quantitative analysis of retinal ganglion cell survival with Rbpms immunolabeling in animal models of optic neuropathies. Invest Ophthalmol Vis Sci 52(13):9694–9702.

Larhammar M, Huntwork-Rodriguez S, Jiang Z, Solanoy H, Sengupta Ghosh A, Wang B, Kaminker JS, Huang K, Eastham-Anderson J, Siu M, Modrusan Z, Farley MM, Tessier-Lavigne M, Lewcock JW, Watkins TA. 2017. Dual leucine zipper kinase-dependent PERK activation contributes to neuronal degeneration following insult. Elife 6.

Lau C, Dolle C, Gossmann TI, Agledal L, Niere M, Ziegler M. 2010. Isoform-specific targeting and interaction domains in human nicotinamide mononucleotide adenylyltransferases. J Biol Chem 285(24):18868–18876.

Lemonidis K, MacLeod R, Baillie GS, Chamberlain LH. 2017. Peptide array based screening reveals a large number of proteins interacting with the ankyrin repeat domain of the zDHHC17 S-acyltransferase. J Biol Chem.

Lemonidis K, Sanchez-Perez MC, Chamberlain LH. 2015. Identification of a Novel Sequence Motif Recognized by the Ankyrin Repeat Domain of zDHHC17/13 S-Acyltransferases. J Biol Chem 290(36):21939–21950.

Lorber B, Tassoni A, Bull ND, Moschos MM, Martin KR. 2012. Retinal ganglion cell survival and axon regeneration in WldS transgenic rats after optic nerve crush and lens injury. BMC Neurosci 13:56.

Mahar M, Cavalli V. 2018. Intrinsic mechanisms of neuronal axon regeneration. Nat Rev Neurosci 19(6):323–337.

Martin DDO, Kanuparthi PS, Holland SM, Sanders SS, Jeong HK, Einarson MB, Jacobson MA, Thomas GM. 2019. Identification of Novel Inhibitors of DLK Palmitoylation and Signaling by High Content Screening. Sci Rep 9(1):3632.

Martin KR, Klein RL, Quigley HA. 2002. Gene delivery to the eye using adeno-associated viral vectors. Methods 28(2):267–275.

Mayer PR, Huang N, Dewey CM, Dries DR, Zhang H, Yu G. 2010. Expression, localization, and biochemical characterization of nicotinamide mononucleotide adenylyltransferase 2. J Biol Chem 285(51):40387–40396.

Milde S, Coleman MP. 2014. Identification of palmitoyltransferase and thioesterase enzymes that control the subcellular localization of axon survival factor nicotinamide mononucleotide adenylyltransferase 2 (NMNAT2). J Biol Chem 289(47):32858–32870.

Milde S, Gilley J, Coleman MP. 2013. Subcellular localization determines the stability and axon protective capacity of axon survival factor Nmnat2. PLoS Biol 11(4):e1001539.

Montersino A, Thomas GM. 2015. Slippery signaling: Palmitoylation-dependent control of neuronal kinase localization and activity. Mol Membr Biol 32(5-8):179–188.

Obenauer JC, Cantley LC, Yaffe MB. 2003. Scansite 2.0: Proteome-wide prediction of cell signaling interactions using short sequence motifs. Nucleic Acids Res 31(13):3635–3641.

Ohno Y, Kihara A, Sano T, Igarashi Y. 2006. Intracellular localization and tissue-specific distribution of human and yeast DHHC cysteine-rich domain-containing proteins. Biochim Biophys Acta 1761(4):474–483.

Palin K, Cunningham C, Forse P, Perry VH, Platt N. 2008. Systemic inflammation switches the inflammatory cytokine profile in CNS Wallerian degeneration. Neurobiol Dis 30(1):19–29.

Roth AF, Feng Y, Chen L, Davis NG. 2002. The yeast DHHC cysteine-rich domain protein Akr1p is a palmitoyl transferase. J Cell Biol 159(1):23–28.

Sajgo S, Ghinia MG, Brooks M, Kretschmer F, Chuang K, Hiriyanna S, Wu Z, Popescu O, Badea TC. 2017. Molecular codes for cell type specification in Brn3 retinal ganglion cells. Proc Natl Acad Sci U S A 114(20):E3974–E3983.

Sanders SS, Parsons MP, Mui KK, Southwell AL, Franciosi S, Cheung D, Waltl S, Raymond LA, Hayden MR. 2016. Sudden death due to paralysis and synaptic and behavioral deficits when Hip14/Zdhhc17 is deleted in adult mice. BMC Biol 14(1):108.

Sasaki Y, Araki T, Milbrandt J. 2006. Stimulation of nicotinamide adenine dinucleotide biosynthetic pathways delays axonal degeneration after axotomy. J Neurosci 26(33):8484–8491.

Sasaki Y, Vohra BP, Baloh RH, Milbrandt J. 2009. Transgenic mice expressing the Nmnat1 protein manifest robust delay in axonal degeneration in vivo. J Neurosci 29(20):6526–6534.

Shin JE, Cho Y, Beirowski B, Milbrandt J, Cavalli V, DiAntonio A. 2012. Dual leucine zipper kinase is required for retrograde injury signaling and axonal regeneration. Neuron 74(6):1015–1022.

Shin JE, Ha H, Kim YK, Cho Y, DiAntonio A. 2019. DLK regulates a distinctive transcriptional regeneration program after peripheral nerve injury. Neurobiol Dis 127:178–192.

Simon DJ, Pitts J, Hertz NT, Yang J, Yamagishi Y, Olsen O, Mark MT, Molina H, Tessier-Lavigne M. 2016. Axon Degeneration Gated by Retrograde Activation of Somatic Pro-apoptotic Signaling. Cell 164(5):1031–1045.

Singaraja RR, Hadano S, Metzler M, Givan S, Wellington CL, Warby S, Yanai A, Gutekunst CA, Leavitt BR, Yi H, Fichter K, Gan L, McCutcheon K, Chopra V, Michel J, Hersch SM, Ikeda JE, Hayden MR. 2002. HIP14, a novel ankyrin domain-containing protein, links huntingtin to intracellular trafficking and endocytosis. Hum Mol Genet 11(23):2815–2828.

Singaraja RR, Huang K, Sanders SS, Milnerwood AJ, Hines R, Lerch JP, Franciosi S, Drisdel RC, Vaid K, Young FB, Doty C, Wan J, Bissada N, Henkelman RM, Green WN, Davis NG, Raymond LA, Hayden MR. 2011. Altered palmitoylation and neuropathological deficits in mice lacking HIP14. Hum Mol Genet 20(20):3899–3909.

Thomas GM, Hayashi T, Chiu SL, Chen CM, Huganir RL. 2012. Palmitoylation by DHHC5/8 targets GRIP1 to dendritic endosomes to regulate AMPA-R trafficking. Neuron 73(3):482–496.

Tortosa E, Adolfs Y, Fukata M, Pasterkamp RJ, Kapitein LC, Hoogenraad CC. 2017. Dynamic Palmitoylation Targets MAP6 to the Axon to Promote Microtubule Stabilization during Neuronal Polarization. Neuron 94(4):809–825 e807.

Tran NM, Shekhar K, Whitney IE, Jacobi A, Benhar I, Hong G, Yan W, Adiconis X, Arnold ME, Lee JM, Levin JZ, Lin D, Wang C, Lieber CM, Regev A, He Z, Sanes JR. 2019. Single-Cell Profiles of Retinal Ganglion Cells Differing in Resilience to Injury Reveal Neuroprotective Genes. Neuron 104(6):1039–1055 e1012.

Verardi R, Kim JS, Ghirlando R, Banerjee A. 2017. Structural Basis for Substrate Recognition by the Ankyrin Repeat Domain of Human DHHC17 Palmitoyltransferase. Structure 25(9):1337–1347 e1336.

Wang JT, Medress ZA, Barres BA. 2012. Axon degeneration: molecular mechanisms of a selfdestruction pathway. J Cell Biol 196(1):7–18.

Wang JT, Medress ZA, Vargas ME, Barres BA. 2015. Local axonal protection by WldS as revealed by conditional regulation of protein stability. Proc Natl Acad Sci U S A 112(33):10093–10100.

Watkins TA, Wang B, Huntwork-Rodriguez S, Yang J, Jiang Z, Eastham-Anderson J, Modrusan Z, Kaminker JS, Tessier-Lavigne M, Lewcock JW. 2013. DLK initiates a transcriptional program that couples apoptotic and regenerative responses to axonal injury. Proc Natl Acad Sci U S A 110(10):4039–4044.

Welsbie DS, Mitchell KL, Jaskula-Ranga V, Sluch VM, Yang Z, Kim J, Buehler E, Patel A, Martin SE, Zhang PW, Ge Y, Duan Y, Fuller J, Kim BJ, Hamed E, Chamling X, Lei L, Fraser IDC, Ronai ZA, Berlinicke CA, Zack DJ. 2017. Enhanced Functional Genomic Screening Identifies Novel Mediators of Dual Leucine Zipper Kinase-Dependent Injury Signaling in Neurons. Neuron 94(6):1142–1154 e1146.

Welsbie DS, Yang Z, Ge Y, Mitchell KL, Zhou X, Martin SE, Berlinicke CA, Hackler L, Jr., Fuller J, Fu J, Cao LH, Han B, Auld D, Xue T, Hirai S, Germain L, Simard-Bisson C, Blouin R, Nguyen JV, Davis CH, Enke RA, Boye SL, Merbs SL, Marsh-Armstrong N, Hauswirth WW, DiAntonio A, Nickells RW, Inglese J, Hanes J, Yau KW, Quigley HA, Zack DJ. 2013. Functional genomic screening identifies dual leucine zipper kinase as a key mediator of retinal ganglion cell death. Proc Natl Acad Sci U S A 110(10):4045–4050.

Yamagishi Y, Tessier-Lavigne, M. 2016. An Atypical SCF-like Ubiquitin Ligase Complex Promotes Wallerian Degeneration through Regulation of Axonal Nmnat2. Cell Rep 17(3):774–782.

Yang J, Wu Z, Renier N, Simon DJ, Uryu K, Park DS, Greer PA, Tournier C, Davis RJ, Tessier-Lavigne M. 2015. Pathological axonal death through a MAPK cascade that triggers a local energy deficit. Cell 160(1-2):161–176.

Young FB, Butland SL, Sanders SS, Sutton LM, Hayden MR. 2012. Putting proteins in their place: palmitoylation in Huntington disease and other neuropsychiatric diseases. Prog Neurobiol 97(2):220–238.

Yue F, Cheng Y, Breschi A, Vierstra J, Wu W, Ryba T, Sandstrom R, Ma Z, Davis C, Pope BD, Shen Y, Pervouchine DD, Djebali S, Thurman RE, Kaul R, Rynes E, Kirilusha A, Marinov GK, Williams BA, Trout D, Amrhein H, Fisher-Aylor K, Antoshechkin I, DeSalvo G, See LH, Fastuca M, Drenkow J, Zaleski C, Dobin A, Prieto P, Lagarde J, Bussotti G, Tanzer A, Denas O, Li K, Bender MA, Zhang M, Byron R, Groudine MT, McCleary D, Pham L, Ye Z, Kuan S, Edsall L, Wu YC, Rasmussen MD, Bansal MS, Kellis M, Keller CA, Morrissey CS, Mishra T, Jain D, Dogan N, Harris RS, Cayting P, Kawli T, Boyle AP, Euskirchen G, Kundaje A, Lin S, Lin Y, Jansen C, Malladi VS, Cline MS, Erickson DT, Kirkup VM, Learned K, Sloan CA, Rosenbloom KR, Lacerda de Sousa B, Beal K, Pignatelli M, Flicek P, Lian J, Kahveci T, Lee D, Kent WJ, Ramalho Santos M, Herrero J, Notredame C, Johnson A, Vong S, Lee K, Bates D, Neri F, Diegel M, Canfield T, Sabo PJ, Wilken MS, Reh TA, Giste E, Shafer A, Kutyavin T, Haugen E, Dunn D, Reynolds AP, Neph S, Humbert R, Hansen RS, De Bruijn M, Selleri L, Rudensky A, Josefowicz S, Samstein R, Eichler EE, Orkin SH, Levasseur D, Papayannopoulou T, Chang KH, Skoultchi A, Gosh S, Disteche C, Treuting P, Wang Y, Weiss MJ, Blobel GA, Cao X, Zhong S, Wang T, Good PJ, Lowdon RF, Adams LB, Zhou XQ, Pazin MJ, Feingold EA, Wold B, Taylor J, Mortazavi A, Weissman SM, Stamatoyannopoulos JA, Snyder MP, Guigo R, Gingeras TR, Gilbert DM, Hardison RC, Beer MA, Ren B. 2014. A comparative encyclopedia of DNA elements in the mouse genome. Nature 515(7527):355–364.

Zeisel A, Hochgerner H, Lonnerberg P, Johnsson A, Memic F, van der Zwan J, Haring M, Braun E, Borm LE, La Manno G, Codeluppi S, Furlan A, Lee K, Skene N, Harris KD, Hjerling-Leffler J, Arenas E, Ernfors P, Marklund U, Linnarsson S. 2018. Molecular Architecture of the Mouse Nervous System. Cell 174(4):999–1014 e1022.

